# Suaeda Salsa Hyperspectral Index (SSHI) for mapping *S. salsa* in coastal wetlands using hyperspectral satellite imagery

**DOI:** 10.1101/2024.12.19.629534

**Authors:** Mengyao Zhang, Yinghai Ke, Kun Shang, Zhaojun Zhuo, Han Liu, Peng Li, Nana Zhao, Jinghan Sha, Jinyuan Li

## Abstract

*Suaeda Salsa (S. salsa),* a pioneer species with short and red-purplish plants in the intertidal zones, has significant ecological, economic, recreational and tourism values. Timely monitoring of *S. salsa* is crucial for understanding its dynamics and sustainable management of coastal wetlands. Hyperspectral satellite offers valuable opportunities due to its detailed spectral information. This study proposed a *Suaeda Salsa* Hyperspectral Index (SSHI) for *S. salsa* mapping based on hyperspectral satellite imagery. The SSHI was developed by considering the large within-class spectral variations of cover types at coastal wetlands, accounting for the spectral correlations in hyperspectral data, and employing dynamic band selection on a per-pixel basis to optimize the separation of *S. salsa* from other land covers. 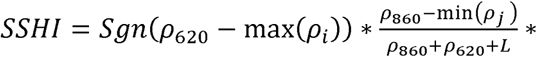 (1+ L), where i∈[499nm, 611nm], j∈[628nm, 851nm], L= 0.5. We applied SSHI on ZY1-02D/E AHSI images over Yellow River Delta (YRD) and Liao River Delta (LRD), China during 2021 to 2023. Based on SSHI, a simple thresholding method and a random forest (RF) model were used to map *S. salsa*. Our results showed that the overall accuracies of *S. salsa* maps achieved 87.96%∼89.43% (YRD) and 92.03%∼93.36% (LRD) using thresholds, and 92.62%∼94.05% (YRD) and 94.74%∼94.79% (LRD) using RF. For RF, incorporating SSHI improves *S. salsa* producer’s accuracies (user’s accuracies) by 1.02%∼6.85% (2.26%∼6.25%) compared to those without SSHI, proving the effectiveness of SSHI. The *S. salsa* maps reveal notable temporal variations, reflecting the impacts of climate change and human activities. SSHI is also applicable to other hyperspectral imagery, such as GF-5B and PRISMA.

## 1. Introduction

Coastal salt marshes play vital roles in shoreline protection, carbon sequestration, biodiversity conservation and strengthening the resilience of coastal ecosystems. *Suaeda salsa* (*S. salsa*) is an important pioneer salt marsh species found in intertidal wetlands in East Asia and Europe [1, 2]. It has short plants (20–60 cm) which normally display red or purple-red tones resulted from increased betacyanin accumulation in response to high salinity and low temperature. In northern coastal China, *S. salsa* communities form beautiful landscape scenery known as "red beaches", which serve as important tourism attractions [3, 4]. However, coastal wetland ecosystems are becoming increasingly fragile due to both natural and human-induced impacts, resulting in the degradation of *S. salsa* [5, 6]. In the past 30 years, the area of *S. salsa* in China has declined by approximately 60% [7], prompting national concern. In response, many regional and local governments have launched restoration projects aimed at *S. salsa* recovery. Timely and accurate monitoring of *S. salsa* is essential to support in-depth research and inform decision-making for restoration initiatives.

While remote sensing techniques have been widely applied in coastal wetlands mapping and vegetation classification, current research on *S. salsa* mapping primarily relies on multispectral satellite imagery [8–12]. Given that *S. salsa* communities exhibit considerable variations in coverage and plant height, and that the soil background in intertidal wetlands presents diverse spectral characteristics due to fluctuating soil moisture and salinity, some studies have reported challenges in accurate mapping of *S. salsa* [12–16]. For instance, Wang et al. (2022) [13] noted considerably lower mapping accuracy for *S. salsa* compared to other vegetation types in the Yellow River Delta when using Landsat 8 OLI data, largely because low-density *S. salsa* was difficult to distinguish from bare flats. In contrast to multispectral data, hyperspectral imagery provides richer spectral detail, enabling more precise differentiation between the spectra of various vegetation species [14, 17, 18]. Furthermore, the distinctive red and purplish tones of *S. salsa* plants present a valuable opportunity for rapid and efficient differentiation of *S. salsa* from other land cover types. Although hyperspectral satellite imagery has been successfully applied to coastal vegetation classification, existing studies often utilized sophisticated classification algorithms and treat *S. salsa* as one of the land cover categories [14, 15, 17].

In this study, we aimed to propose a novel hyperspectral index for the accurate mapping of *S. salsa*, termed *S. Salsa* Hyperspectral Index (SSHI). This index was developed by comprehensively accounting for the spectral variations in coverages of different vegetation types and in soil backgrounds in coastal wetlands. Unlike most existing studies that visually select feature wavebands for constructing hyperspectral indices [17–19], our approach determined the optimal band combination through spectral correlation analysis, optimum index factor (OIF) and Jeffreys-Matusita (J-M) distance. This approach not only enhances the separation of *S. salsa* from other land cover types but also mitigates spectral redundancy commonly present in hyperspectral data. We then applied the SSHI on five ZY1-02D and ZY1-02E AHSI image scenes covering two representative river deltas in China, namely Yellow River Delta (YRD) and Liao River Delta (LRD), for the period between 2021 and 2023. To validate the efficacy of SSHI, we first compared it with other hyperspectral indices associated with red-tone leaf pigments, focusing on its ability to differentiate *S. salsa* from other land cover types. Subsequently, we employed both threshold-based and random forest classification methods to identify *S. salsa* from ZY1-02D/E images. Finally, we generated *S. salsa* distribution maps for different years and conducted a comparative analysis to assess the dynamics of *S. salsa* in both study areas. The applicability of SSHI on other hyperspectral satellite imagery was also discussed.

## 2. Materials

### 2.1 Study area

Two representative coastal wetlands in Northern China were selected for this study: YRD and LRD (Figure 1).

**Figure 1.**
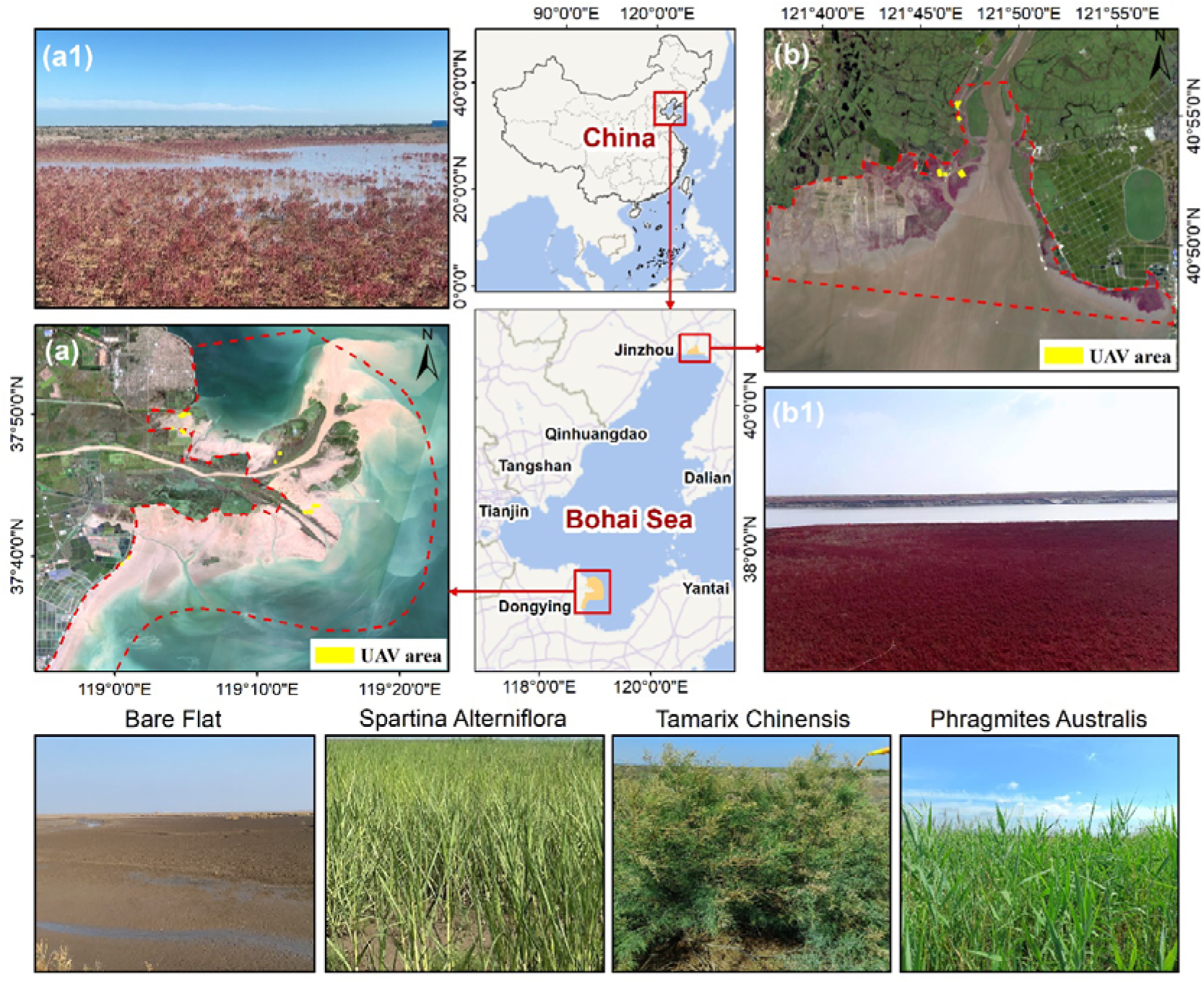
Location and boundaries of the study sites in the North China river deltas: (a) YRD (September 21, 2021, ZY1-02D AHSI), and (b) LRD (September 6, 2022, ZY1-02D AHSI). Subfigures (a1) and (b1) are field photos showing *S. salsa* in YRD and LRD, respectively. The bottom section includes field photos of other land cover types in the study areas.

The YRD study area is within the Yellow River Delta National Nature Reserve (YRDNNR) in Dongying City, Shandong Province, China (118°33′E-119°20′E, 37°35′N-38°12′N), situated in the southern Bohai Bay. It has a temperate continental monsoon climate, with average annual precipitation of about 550 mm and an average temperature of about 12c. Dominant vegetation species include *S*.

*salsa*, *Phragmites australis* (*P. australis*), *Tamarix chinensis* (*T. chinensis*) and the invasive species *Spartina alterniflora* (*S. alterniflora*). Over the past 35 years, *S. salsa*’s habitat has declined by nearly 70% due to the rapid expansion of *S. alterniflora* and human activities [20, 21]. Since 2019, YRDNNR has implemented ecological restoration projects to recover *S. salsa* marshes using both natural regeneration and artificial seeding methods [22]. Furthermore, a large-scale *S. alterniflora* removal project initiated in 2021 has eradicated over 90% of this invasive species by the end of 2023, resulting in significant landscape transformations [23].

The LRD study area, situated in Panjin City, Liaoning Province, China (121°28’E -121°58’E, 40°45’N∼41°06’N), is located on the northern Bohai Sea coast. It has a warm temperate continental semi-humid monsoon climate, with an average annual precipitation of about 650 mm and a mean temperature of 8.3L. Dominant vegetation species are *P. australis* and *S. salsa*. Compared to YRD, *S. salsa* in LRD grows more densely. In recent years, the famous “Red Beach” in LRD has rapidly degraded due to climate change and human impacts, and *S. salsa* has gradually been replaced by *P. australis* and bare flat [24]. Since 2015, large-scale restoration initiatives, including replanting *S. salsa,* dredging tidal channels, and reclaiming intertidal wetlands, have been conducted to conserve the coastal ecosystem. In 2019, Panjin City launched a large-scale project to restore *S. salsa*, creating suitable habitats for its growth. Currently, these restoration efforts are ongoing [25, 26].

### 2.2 Data

#### 2.2.1 ZY1-02D/E hyperspectral satellite images and preprocessing

In this study, we used the hyperspectral images collected by ZY1-02D and ZY1-02E satellites for SSHI development and *S. salsa* mapping. ZY1-02D, a Chinese Earth observation satellite, was launched on September 12, 2019. It is equipped with an 8-band visible and near-infrared camera (VNIC) and a 166-band advanced hyperspectral imager (AHSI). The ZY1-02D AHSI comprises 76 visible near-infrared (VNIR) bands with a 10nm spectral resolution from 400nm to 1040nm and 90 short-wave infrared (SWIR) bands with a 20nm spectral resolution from 1005nm to 2500nm. The satellite offers a spatial resolution of 30m, a quantization level of 12bit, and swath width of 60km (Table 1). ZY1-02E, launched on December 26, 2021, inherits the VNIC and AHSI from ZY1-02D and incorporates an additional long-wave infrared camera. AHSI images of the Yellow River Delta (YRD) were acquired in September 2020, 2021, and 2023, while those of the Liao River Delta (LRD) were captured in September 2022 and August 2023 (Table 1).

**Table 1.** Satellite images and the acquisition date. YRD: Yellow River Delta; LRD: Liao River Delta; AHSI: Advanced Hyperspectral Imager.

| Study area | Hyperspectral satellite image | Date | High-spatial-resolution satellite image | Date |
| --- | --- | --- | --- | --- |
| YRD | ZY1-02D AHSI | 2020-09-26 | Jilin1-KF01A | 2020-08-09 |
|  | ZY1-02D AHSI | 2021-09-29 | Jilin1-KF01A | 2021-09-22 |
|  | ZY1-02E AHSI | 2023-09-07 | Jilin1-GF02D | 2023-09-15 |
| LRD | ZY1-02D AHSI | 2022-09-06 | PlanetScope Dove-R | 2022-09-11 |
|  | ZY1-02E AHSI | 2023-08-09 | PlanetScope Dove-R | 2023-08-09 |

For each of the ZY1-02D/E imagery, the overlapping bands between VNIR bands and SWIR bands were removed, and the bands with the center wavelengths of 1358 nm to1425 nm, 1812 nm to 1947 nm, and 2501 nm were removed because of missing data and stripping problem. Radiometric calibration, FLASHH atmospheric correction and geometric correction were then performed with ENVI software. Savitzky-Golay (S-G) filtering was applied to smooth the spectral curves while preserving important spectral features.

#### 2.2.2 High spatial resolution satellite images

To collect reference samples, we acquired high-spatial-resolution Jilin-1 satellite images and PlanetScope images. Jilin-1 satellites are equipped with sub-meter wide-area cameras that capture 0.75-meter panchromatic images and 3-meter multispectral images across blue, green, red, and near-infrared bands. PlanetScope satellites provide 3-meter resolution, 4-band multispectral (blue, green, red and NIR) imagery. Cloud-free images temporally aligned with the ZY1-02D/E image acquisitions were selected (Table 1). Preprocessing of Jilin-1 images included radiometric calibration, FLAASH atmospheric correction, geometric correction, and pan-sharpening, yielding 4-band multispectral images with a 0.75-meter spatial resolution. For PlanetScope data, Level 3B surface reflectance products were downloaded directly from the Planet website (https://www.planet.com/), eliminating the need for further preprocessing.

#### 2.2.3 Field surveys and Unmanned Aerial Vehicle images

Several field surveys were conducted in August and September during 2020 to 2023 at YRD and LRD, during which researchers located and recorded land cover and vegetation types using high-precision hand-held GPS units. In the field surveys between 10:00 and 14:00, Unmanned Aerial Vehicle (UAV) images were captured over *S. salsa* marshes by a DJI Phantom 4 Multispectral (P4M) drone, which integrates a 6-camera RGB and multispectral imaging system to provide center-meter level images. A total of 15 UAV scenes over YRD and 6 scenes over LRD with average area of 9.5 hectares and spatial resolution of 2.64 cm (flight height of 50 m) were collected. DJI Terra synchronization processing software was used for preprocessing, including image stitching and radiometric correction, to obtain UAV multispectral image maps of the study area. Both field survey records and the UAV imagery were used to assist collection of reference samples (Section 3.1.1 and Section 3.2).

## 3. Methods

### 3.1 Development of *S. salsa* hyperspectral index (SSHI)

In this study, we first obtained and analyzed the hyperspectral curves of various land cover and vegetation types, accounting for variations in vegetation coverage and soil moisture (Section 3.1.1 and 3.1.2). Based on this analysis, we identified the optimal band combination that maximized information content, minimized redundancy, and was most effective for distinguishing *S. salsa* (Section 3.1.3). Using these selected bands, we developed the SSHI (Section 3.1.4), which was subsequently applied to the YRD and LRD to map the distribution of *S. salsa* and analyze its dynamics.

#### 3.1.1 Selection of sample datasets for SSHI development

The land cover types in the intertidal wetlands in northern coastal China can be generally categorized into: water, bare flat, and saltmarsh vegetation which primarily include *S. salsa*, *S. alterniflora*, *P. australis*, and *T. chinensis*. In this study, we grouped *S. alterniflora*, *P. australis*, and *T. chinensis* into “green vegetation” because they have similar spectral characteristics [14]. Therefore, we examined and compared the spectral curves of *S. salsa*, green vegetation, bare flat, and water. Considering the large within-class spectral variations, we further examined the spectral curves of bare flat samples representing different levels of soil moisture, as well as those of *S. salsa* and green vegetation samples representing different levels of fractional coverages.

First, we visually delineated the approximate extent of bare flat from the Jilin-1 imagery acquired on September 22, 2021, and within this extent, we calculated the Normalized Differential Water Index from ZY1-02D imagery on September 29, 2021 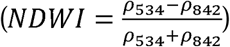 [27, 28]. Although field-based soil moisture measurements are not available, NDWI have proved to be a reliable and widely-used indicator of water content levels [29–32]. Then, the bare flat NDWI (ranging from -0.4 ∼ -0.1) was grouped into three levels based on the quantiles: Level 1 (NDWI ≤ -0.25), Level 2 (-0.25 < NDWI ≤ -0.18), and Level 3 (NDWI > -0.18), which represent low-, mid- and high-water content, respectively. From each level, we randomly selected 200 pixels, totaling 600 bare flat samples.

Next, we visually delineated the approximate extent of vegetation from the Jilin-1 imagery. Green vegetation patches can be easily delineated by visual interpretation, while it could be challenging to accurately delineate low-density *S. salsa* patches because of the short plant heights. In our study, we utilized the UAV imagery and the field records as the reference to assist visual interpretation (Table

S1). Within the green vegetation and *S. salsa* patches, we calculated the NDVI from the ZY1-02D imagery 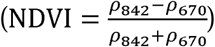, which has been widely recognized as an indicator representing fractional coverage of vegetation. We then respectively grouped the NDVI ranges of *S. salsa* patches and green vegetation patches into four levels. For *S. salsa*, the four levels are: Level 1 (0.12 ≤ NDVI < 0.21), Level 2 (0.21≤ NDVI <0.32), Level 3 (0.33 ≤ NDVI < 0.44), Level 4 (NDVI ≥ 0.44). For green vegetation the four levels are: Level 1 (0.12 ≤ NDVI < 0.26), Level 2 (0.26≤ NDVI <0.44), Level 3 (0.44 ≤ NDVI < 0.59), Level 4 (NDVI ≥ 0.59) (examples shown in Table S1). For each NDVI level, we randomly selected 125 pixels, totaling 600 *S. salsa* samples and 600 green vegetation samples. We also randomly selected 300 water samples from the ZY-01D imagery. To sum up, a total of 2100 samples including 300 water samples, 600 bare flat samples with three soil moisture levels, 600 *S. salsa* samples and 600 green vegetation samples with four coverage levels were generated (Figure S1). The spectral curves of these samples were analyzed and compared (Figure 2).

**Figure 2.**
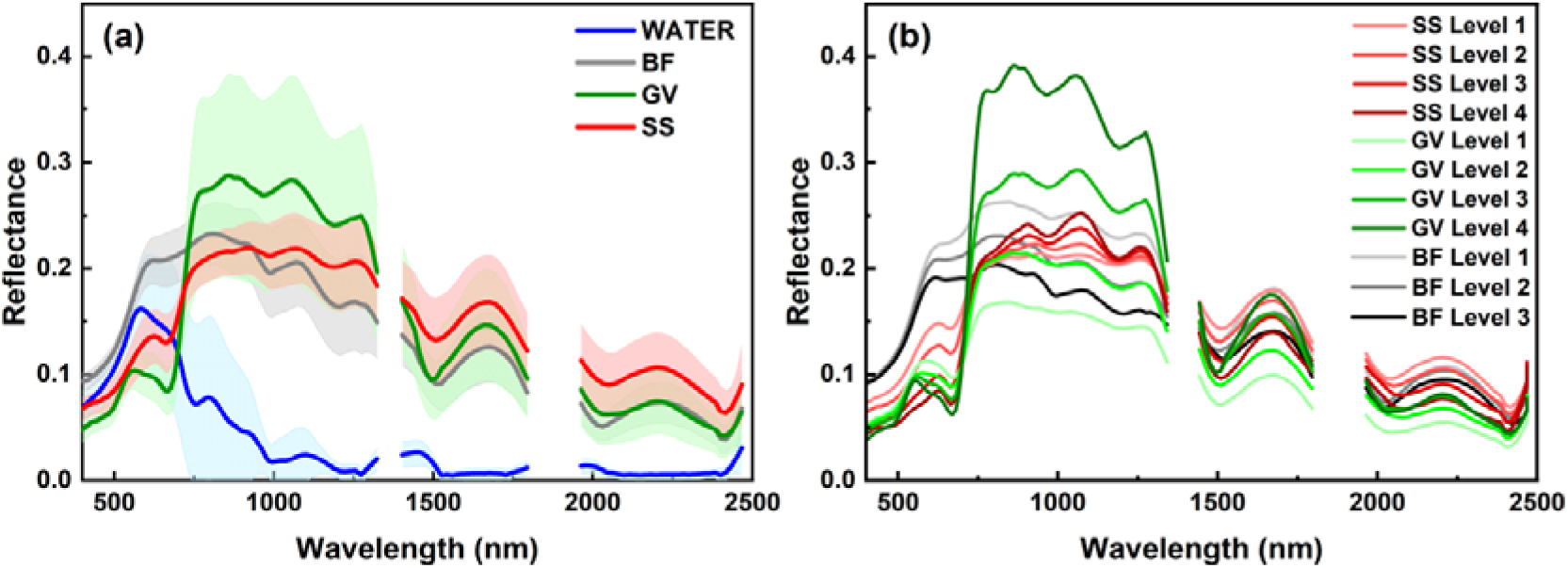
Hyperspectral curves of land cover types. (a) Average spectral curves and reflectance ranges (Mean ± SD) of four land cover types; (b) Average spectral curves of *S. salsa* (SS) and green vegetation (GV) with different NDVI levels, and bare flat (BF) with different NDWI levels.

#### 3.1.2 Analysis of spectral characteristics

The spectral curves of the selected samples (Section 3.1.1) exhibit significant within-class variations (Figure 2). Water shows distinct spectral characteristics compared to bare flats and vegetation, with much lower reflectance in the NIR wavebands than in the visible bands (Figure 2a). Both *S. salsa* and green vegetation have notably higher NIR reflectance compared to the visible bands; however, the key distinction lies in their visible wavelength responses: *S. salsa* peaks in the red wavelengths, whereas green vegetation peaks in the green wavelengths (around 499–576 nm) (Figure 2a). Bare flats also display higher NIR reflectance than visible bands, but their visible-band reflectance exceeds that of *S. salsa* and green vegetation. Furthermore, drier bare flats exhibit higher red-band and NIR reflectance compared to wet bare flats (Figure 2b). As vegetation coverage decreases, *S. salsa* shows increased visible-band reflectance and reduced NIR reflectance. Similarly, low-coverage green vegetation exhibits higher green/red reflectance and lower NIR reflectance compared to high-coverage vegetation, with the green peak shifting to longer wavelengths as vegetation coverage decreases.

#### 3.1.3 Determination of optimal band combination

Hyperspectral images contain a large number of spectral bands with strong reflectance correlation between neighboring bands. To avoid data redundancy, we selected the optimal band combination using correlation analysis, optimum index factor (OIF), and Jeffreys-Matusita (J-M) distance analysis [33].

First, we calculated the correlation coefficient to evaluate information overlap among the bands. Figure 3 shows the correlation coefficient matrix for each band pair, excluding pixels with NDVI < 0 to focus solely on land pixels. The matrix reveals three highly correlated subspaces: 400–722 nm (subspace 1), 730–1308 nm (subspace 2), and 1324–2450 nm (subspace 3), with average correlation coefficients of 0.994, 0.990, and 0.976, respectively. This indicates substantial information redundancy among bands within each subspace. To minimize redundancy, bands from different subspaces were prioritized for selection. Considering that the primary distinction of *S. salsa* lies in its peak reflectance within the red wavelength range (Figure 2), specifically at the center wavelength of 620 nm, the 620 nm band was chosen from the first subspace.

**Figure 3.**
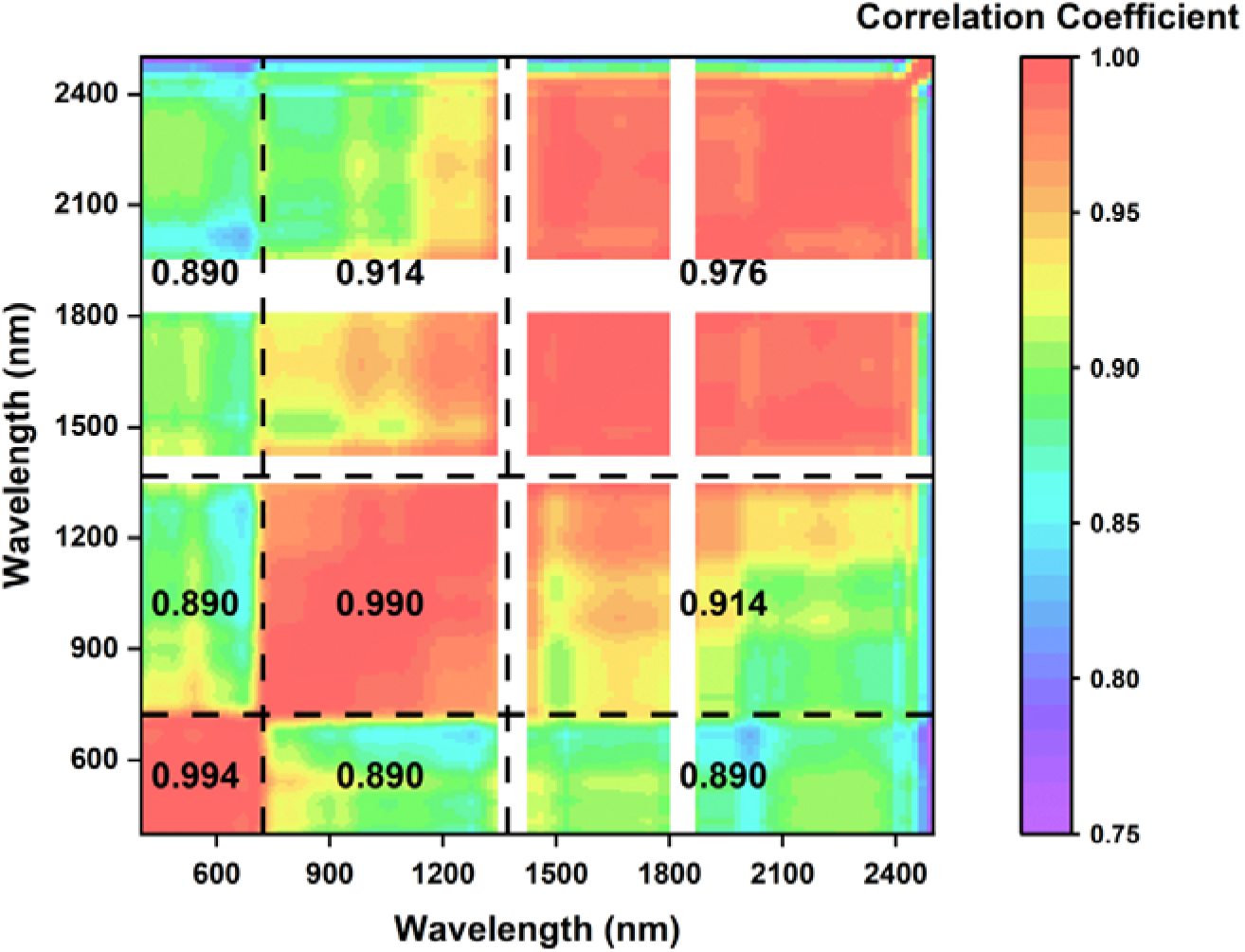
Band correlation matrix.

To select bands from subspaces 2 and 3, we calculated OIF between the band 620nm and each band in the other subspaces. OIF is a common indicator for optimal band combination selection, calculated by the standard deviation and correlation between bands (Equation1). Higher OIF values

indicate more effective information and less redundancy [34, 35].

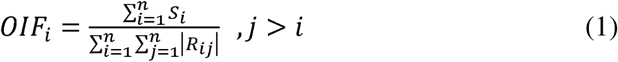

where S_i_ is the standard deviation of band i reflectance, R_ij_ is the correlation coefficient between the reflectance of band i and j, and n is the number of bands. We calculated OIF values of two-band and three-band combinations containing 620nm. Table 2 lists the highest ten OIF values of these combinations. For the two-band combination, the OIF values between 620nm and the bands in subspace 2 are higher than the other two-band combination between 620nm and subspace 3, and also higher than the three-band combinations, indicating that combination of 620nm and subspace 2 provide more complementary information. Therefore, the top ten combinations of 620nm and the bands in subspace 2 were selected as the candidate band combinations.

**Table 2.**
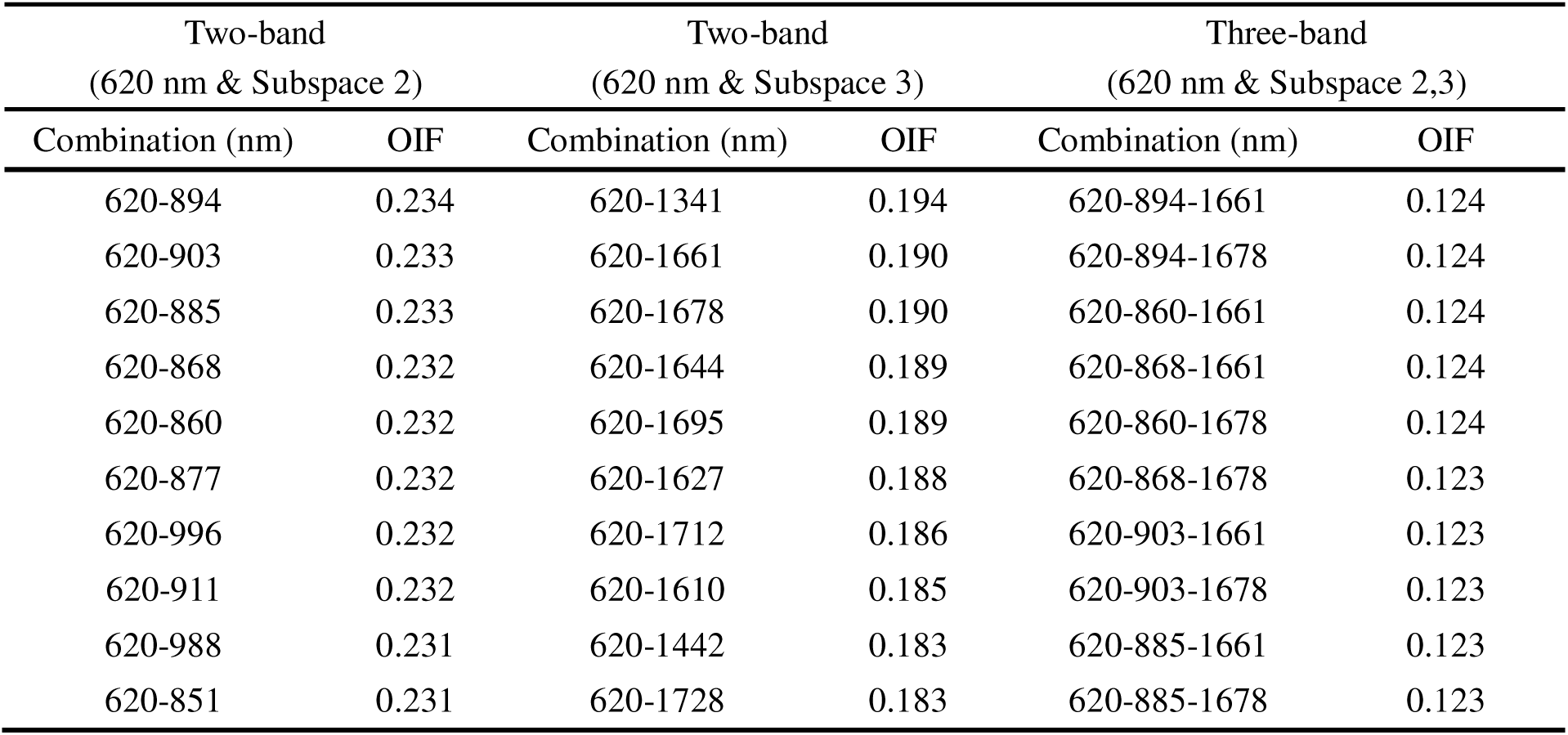
OIF of different band combinations.

From these candidates, we calculated the interclass separability between *S. salsa* and other land cover types using J-M distance based on the samples [36, 37]. The combination with the highest J-M distance value, 620 nm and 860 nm, was selected as the optimal band combination for SSHI development (Table 3)

**Table 3.**
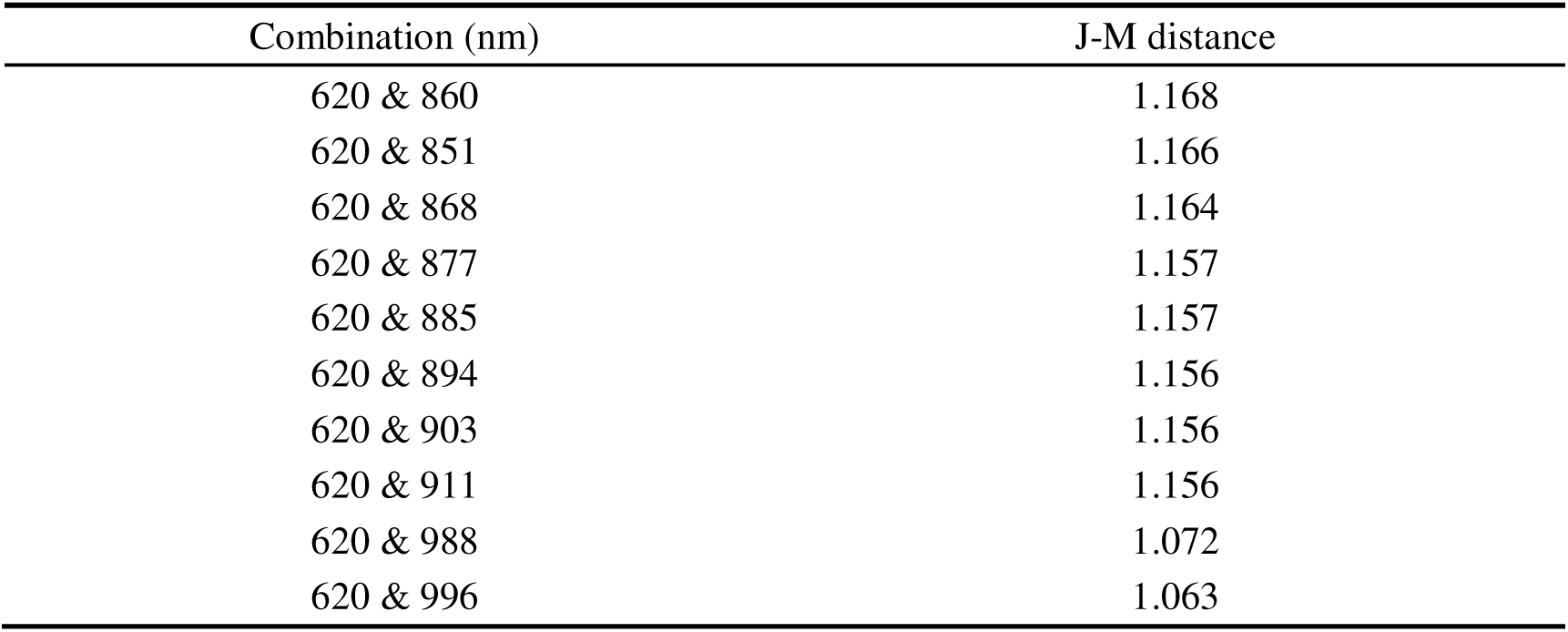
J-M distance of different band combinations.

#### 3.1.4 Formulation of SSHI

The objective of the SSHI is to enhance the distinguishment between *S. salsa* and other land cover types. As water is easily distinguishable, in this study we focus on the distinction of *S. salsa* from green vegetation and bare flats. The primary goal of the SSHI was to enhance the differentiation between *S. salsa* and other land cover types. Since water is easily distinguishable, in this study we focus on separating *S. salsa* from green vegetation and bare flats. Figure 4 illustrates that *S. salsa*, regardless of its fractional coverage, exhibits a local reflectance maximum at 620 nm within the visible spectrum. In contrast, green vegetation shows its local reflectance maximum in the green band, which shifts towards 620 nm as coverage decreases. This shift is attributed to the higher red reflectance of bare flats compared to the green band. To address this, we employed a “max” function to dynamically select the maximum reflectance value within the center wavelength range of 499–611 nm on a pixel-by-pixel

**Figure 4.**
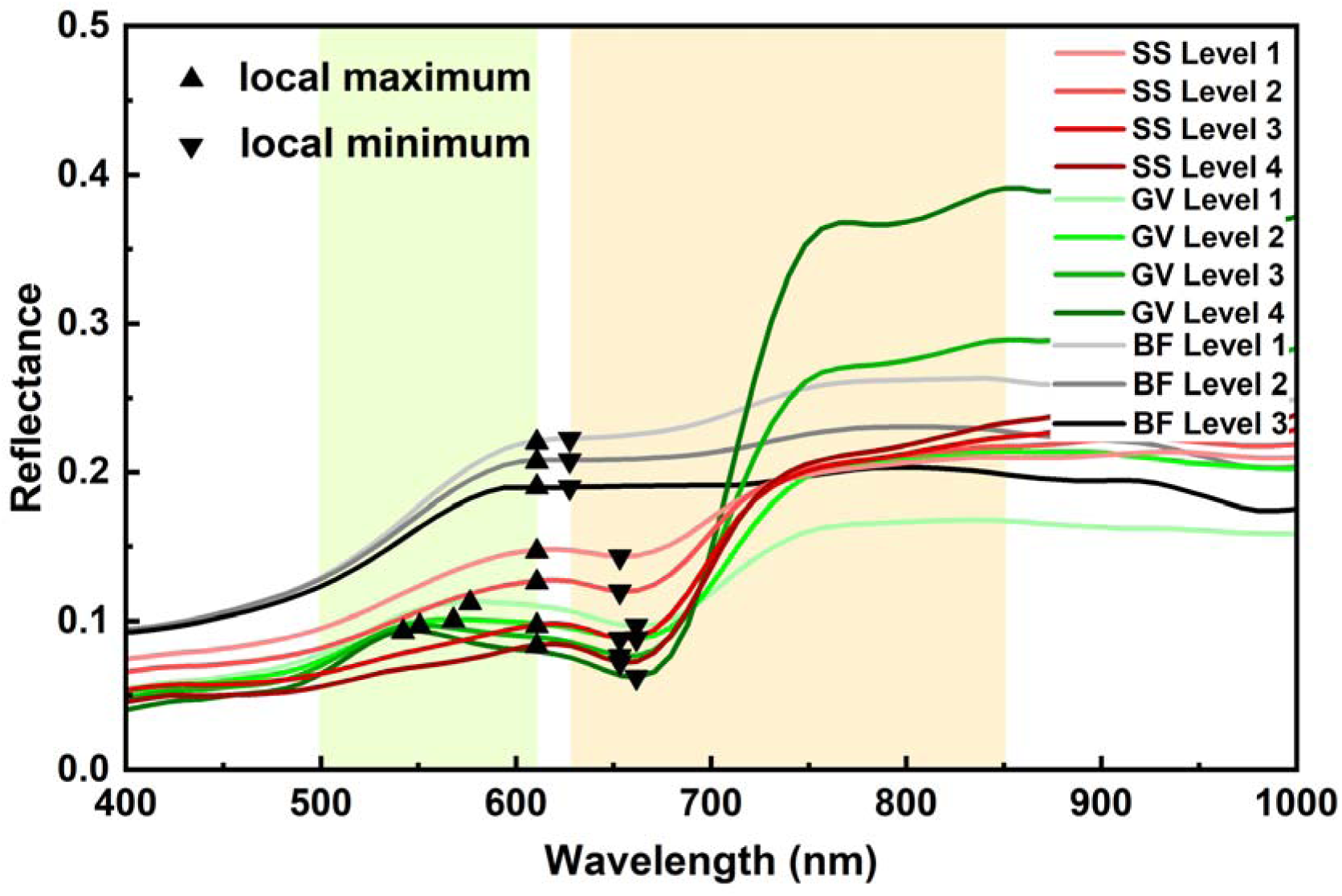
Detailed illustration of spectral curves in the range of 400 nm to 1000 nm. Green shade represents the wavelength range 499 nm∼611 nm, orange shade represents the wavelength range 628 nm∼851 nm.

basis (Figure 4) and compared it to the reflectance at 620 nm (p_620_) (Equation 2).

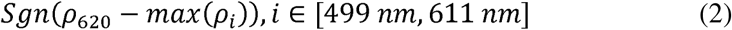

where max(p_i_) denotes the maximum reflectance within the range from 499nm to 611nm. Negative values of (p_620_ - max(p_i_)), i.e., Sgn(p_620_ - max(p_i_)) = -1, suggest green vegetation, while positive values suggest *S. salsa* or bare flat.

A comparison of the spectral curves of *S. salsa* and bare flats (Figure 4) reveals that *S. salsa* exhibits relatively lower reflectance in the visible bands. Notably, *S. salsa* shows a local minimum in reflectance between the center wavelengths of 628 nm and 851 nm, followed by a rapid increase from the red edge to the NIR range. In contrast, bare flats display a much slower increase in reflectance from the visible to NIR wavelengths. To distinguish *S. salsa* from bare flats, we adopted a vegetation index approach. Specifically, the Soil Adjusted Vegetation Index (SAVI), given by 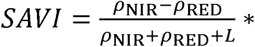 (1 + L), was chosen for its proven accuracy in monitoring vegetation with low coverage [38]. To further enhance the distinction between NIR and red reflectance, we modified the SAVI numerator by substituting the red band reflectance with the minimum reflectance value within the center wavelength range of 628–851 nm (Equation 3).

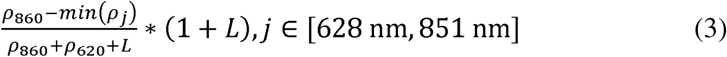

where min(p_j_) is the minimum value of reflectance between the central wavelength of 628nm to 851nm; L = 0.5, which can minimize the effect of soil background reflection [38].

The *Suaeda Salsa* Hyperspectral Index (SSHI) (Equation 4) was formulated as the product of

Equation 2 and Equation 3:

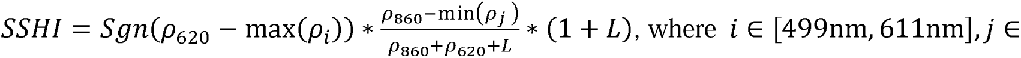

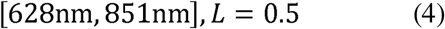

#### 3.2 Mapping *S. salsa* based on SSHI

First, all pixels with NDVI < 0 were masked out, as vegetation cannot have a negative NDVI. Subsequently, we calculated the SSHI and employed both an empirical threshold-based method and a random forest model to classify *S. salsa* and non-*S. salsa* areas. Since both approaches are supervised, we generated training and test samples for each hyperspectral image (Table 1) through visual interpretation of Jilin-1 or PlanetScope high-resolution imagery acquired in the same year (Section 2.2.2), along with UAV imagery and field survey records. UAV imagery was particularly useful for assisting the manual interpretation of low-coverage *S. salsa* samples. Random points were generated, and their cover types (*S. salsa*, green vegetation, bare flats) were determined by three remote sensing experts. Uncertain samples were excluded to ensure data quality, and a minimum of 500 samples was retained for each class. For each image, a total of 1600 samples were prepared, including 500 *S. salsa* samples and 1100 non-*S. salsa* samples. Of these, 1000 samples were randomly selected for training, while the remaining 600 were reserved for validation.

#### 3.2.1 Threshold-based method

The comparison of SSHI of *S. salsa* and other land cover types found that *S. salsa* generally have SSHI over 0.2, while bare flat has low SSHI values and green vegetation has negative SSHI (shown in Figure 5). Therefore, for each image we tried SSHI thresholds within the range from 0.1 to 0.3 with 0.01 step size. The SSHI value producing the highest overall accuracies (Section 3.3.2) that were evaluated with the training datasets were determined as the optimal SSHI threshold. The optimal threshold was then used to binarize the image into *S. salsa* and non-*S. salsa*.

**Figure 5.**
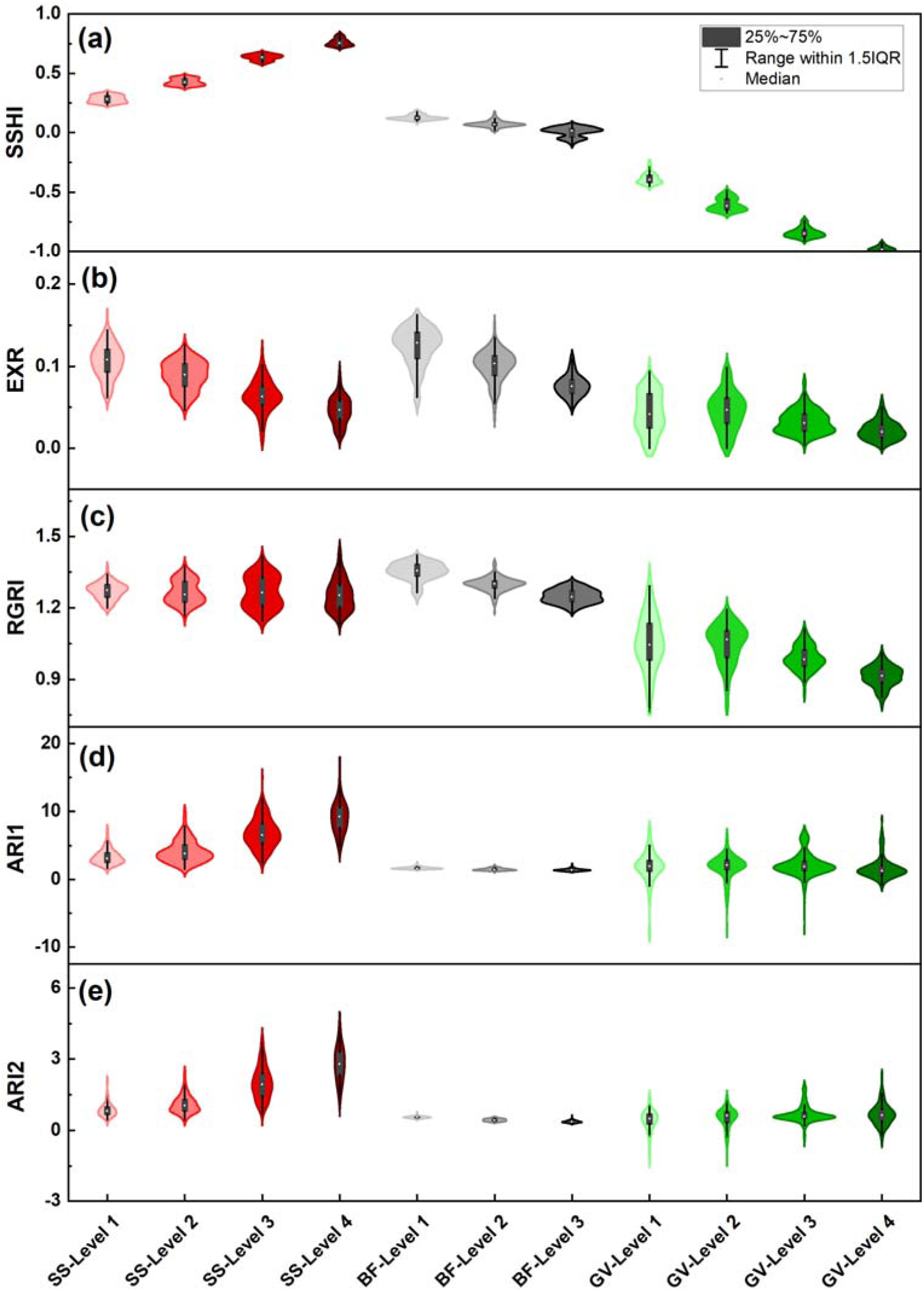
Spectral index values of the *S. salsa* (SS), green vegetation (GV), and bare flat (BF) samples. SS-Level1 to SS-Level4 (GV-Level1 to GV-Level4) denote the *S. salsa* (green vegetation) samples with low to high coverages (represented as NDVI levels); BF-Level1 to BF-Level 3 denote the bare flat samples with low to high soil water content (represented as NDWI levels).

#### 3.2.2 Random Forest classification

Random forest (RF) algorithm is a supervised machine learning technique that has been widely used for remote sensing image classification [39]. In this study, we selected 35 wavebands within 391-598nm, 658-709nm, 881-898nm, 967-984nm, 1001-1044nm, and 1972-2002nm as classification features because previous studies have reported the importance of these bands in coastal vegetation classification [10, 14, 40]. To assess the performance of SSHI, we designed two classification schemes:

1. RF with SSHI and surface reflectance at the 35 spectral bands, and (2) RF with only the 35 spectral bands. To further evaluate the contribution of SSHI in RF classification, SHapley Additive exPlanations (SHAP) values [41] were calculated to representing each feature’s average marginal contribution to the results.

### 3.3 Assessment of SSHI

#### 3.3.1 Comparison with other hyperspectral indices

To examine the effectiveness of SSHI, we compared it with other hyperspectral indices related to vegetation red pigments including anthocyanins and betacyanins [42], including Excess Red Index 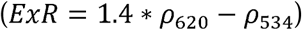 [43] and Red Green Ratio Index 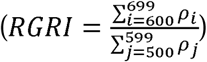 [44], Anthocyanin Reflectance Index 1 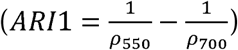 [45], and Anthocyanin Reflectance Index 2 (ARI2 =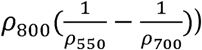) [45]. ExR specifically focuses on the red wavelength, and RGRI has been used to retrieve betacyanin and anthocyanins of *S. salsa* [4]. ARI1 and ARI2 are both sensitive to anthocyanins content [46].

We first calculated the ExR, RGRI, ARI1 and ARI2 for the *S. salsa*, green vegetation and bare flat samples generated in Section 3.1.1, and visually compared the histograms. Then, we calculated the separability index (SI) [47] of the four indices between *S. salsa* and non-*S. salsa* on each of the fiveimages based on the training and validation samples (Section 3.2). SI defines the separability between a pair of classes as the ratio of the inter-class and the intra-class variability:

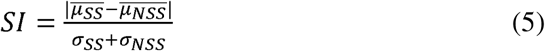

where 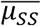 and 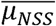 denote the mean index values of *S. salsa* and non-*S. salsa*, respectively; a_ss_ and a_Nss_ denote the corresponding standard deviation of the index values.

#### 3.3.2 Accuracy assessment of mapping results

All *S. salsa* maps generated from the threshold-based and random forest classification were evaluated using the accuracy metrics including Overall Accuracy (OA), Producer’s Accuracy (PA), User’s accuracy (UA) and Kappa coefficients. Detailed formulas are referred to Congalton (1988) [48].

## 4. Results

### 4.1 Comparison with other hyperspectral indices

Figure 5 illustrates the SSHI, ExR, RGRI, ARI1 and ARI2 values of the samples at YRD (Section 3.1.1). Figure 5a shows a clear separation of SSHI between *S. salsa* and both green vegetation and bare flats. All green vegetation samples have negative SSHI values, while *S. salsa* samples exhibit positive values. Bare flats have SSHI values near zero. It appears that SSHI over the threshold around 0.25 could distinguish *S. salsa* from other types. As the coverage of *S. salsa* increases, its SSHI value rises, while the SSHI value for green vegetation decreases with increasing coverage.

Both ExR and RGRI show large overlaps among the three cover types, indicating that the EXR and RGRI can hardly separate *S. salsa* from bare flat and green vegetation (Figure 5b and 5c). For ARI1 and ARI2, bare flat samples show very narrow value range, indicating that both indices are not sensitive to soil water content. However, bare flats still show slight overlaps with low-coverage *S. salsa*. Likely, ARI1 and ARI2 values of green vegetation also illustrates overlaps with those of *S. salsa* (Figure 5d and 5e). Across all five images over YRD and LRD, SSHI consistently demonstrates a much higher separability between *S. salsa* and non-*S. salsa* compared to the other four indices (Figure 6), highlighting its superior performance over the existing red-pigment-related spectral indices for identifying *S. salsa*.

**Figure 6.**
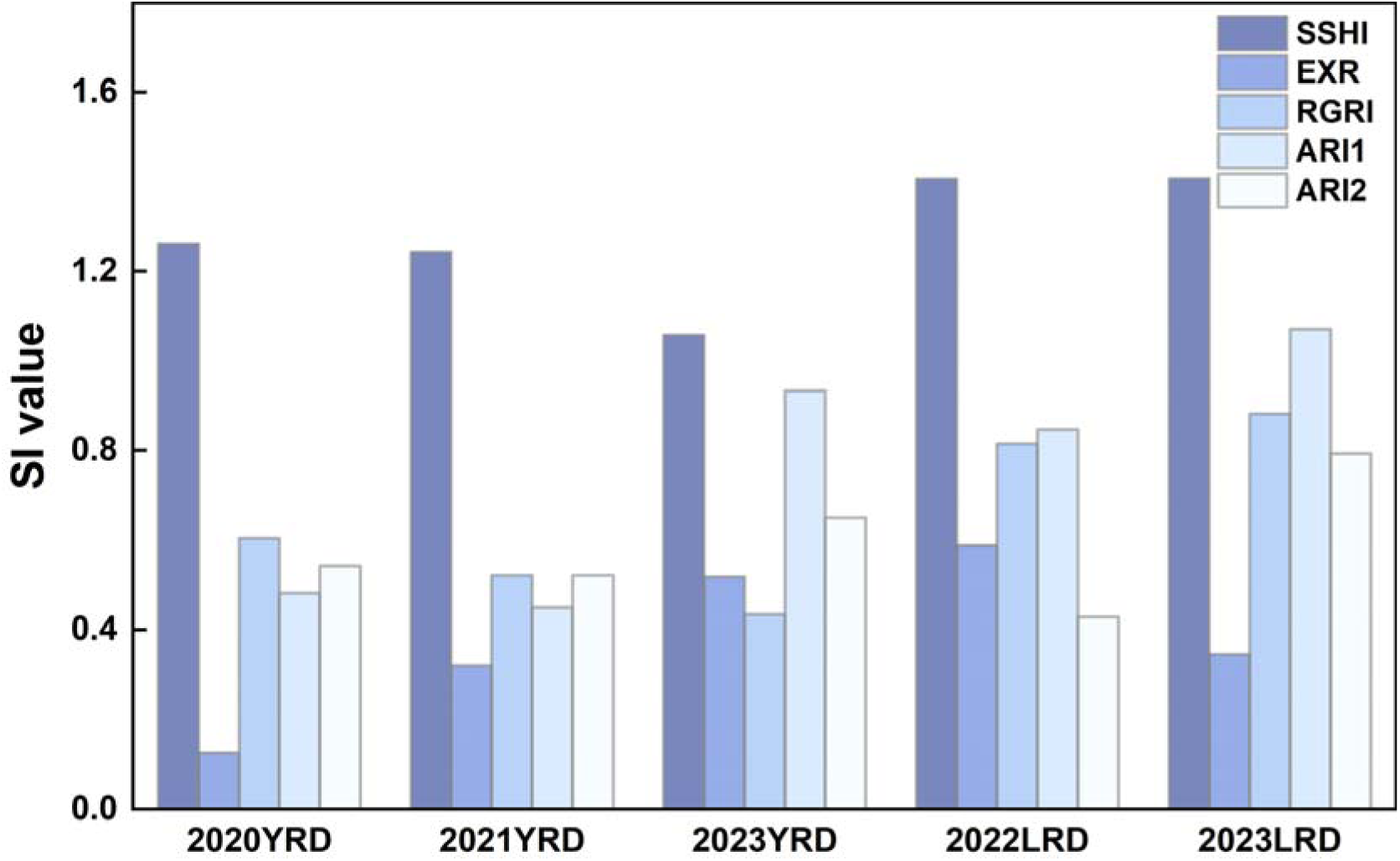
Separability Index values of SSHI, EXR, RGRI, ARI1 and ARI2 indices between *S. salsa* and non-*S. salsa*.

### 4.2 Accuracy assessment of threshold-based *S. salsa* identification

The *S. salsa* maps were generated using both thresholding method and random forest classification models. Figure 7a shows that the optimal thresholds for YRD are 0.22, 0.24 and 0.25 for the year 2020, 2021 and 2023, respectively. Pixels with SSHI greater than the thresholds were identified as *S. salsa*. These thresholds achieved overall accuracies of 89.45%, 89.30% and 90.38% for the training samples. With these thresholds, the evaluation based on the validation samples resulted in OAs of 89.12%, 87.96% and 89.43% (Table 4), which are close to the training accuracies. The optimal thresholds for LRD are 0.20 for both 2022 and 2023, resulting in OAs of 93.43% and 93.39% (Figure 7b). The validation accuracies in LRD are also close to the training accuracies, with OAs of 93.36% and 92.03% for 2022 and 2023, respectively (Table 4). The accuracies in LRD are higher than those in YRD. For YRD, the class of *S. salsa* had both PAs and UAs around 80.00% ∼ 90.26%, and for LRD, the PAs and UAs of *S. salsa* are 86.07% ∼ 95.05%.

**Figure 7.**
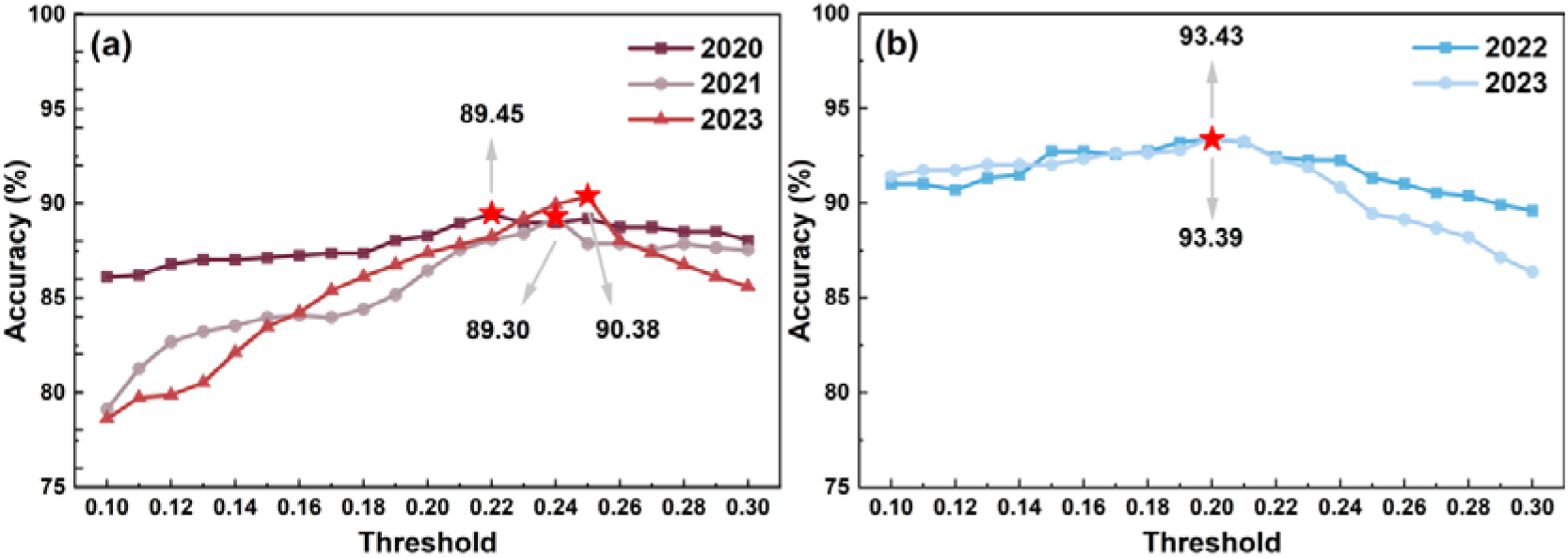
Analysis results of optimal threshold accuracy for (a) YRD and (b) LRD.

**Table 4.** Accuracy metrics of the *S. salsa* maps from optimal threshold method (SS: *S. salsa*; NSS: non-*S. salsa*; OA: overall accuracy; PA: producer’s accuracy; UA: user’s accuracy; Kappa: Kappa coefficient).

| Region/Year | PA (%) |  | UA (%) |  | Kappa | OA (%) |
| --- | --- | --- | --- | --- | --- | --- |
|  | SS | NSS | SS | NSS |  |  |
| YRD 2020 | 80.40 | 93.68 | 86.96 | 90.13 | 0.77 | **89.12** |
| YRD 2021 | 81.62 | 90.91 | 80.75 | 91.37 | 0.72 | **87.96** |
| YRD 2023 | 90.26 | 89.03 | 80.00 | 94.95 | 0.77 | **89.43** |
| LRD 2022 | 95.05 | 92.53 | 86.07 | 97.47 | 0.87 | **93.36** |
| LRD 2023 | 90.56 | 92.93 | 88.59 | 94.20 | 0.84 | **92.03** |

### 4.3 Accuracy assessment of RF-based *S. salsa* identification

Table 5 shows that the RF models based on the combination of SSHI and spectral reflectance yield high accuracies of *S. salsa* maps, with OA of 92.62% ∼94.05% in YRD and 94.74%∼94.79% in LRD. The OAs, PAs (86.32%∼93.41%) and UAs (86.83%∼94.29%) of *S. salsa* are significantly higher than those based solely on surface reflectance (without SSHI). Compared to the classification schemes without SSHI, the incorporating SSHI considerably improved the PAs and UAs of *S. salsa* in both areas, with a more notable improvement in YRD. With SSHI, the PAs increased by 1.02% ∼ 6.85%, and the UAs increased by 4.59% ∼6.25% in YRD; the PAs and UAs in LRD increased by 1.11%∼3.73%. The OA in LRD was higher than that in YRD, exceeding 94% in both 2021 and 2023. In YRD, the UA of *S. salsa* was highest in 2020 (92.75%), but dropped to 90.61% in 2021 and 86.83% in 2023. The PA of *S. salsa* was highest in 2023 (91.28%), while in 2020 and 2021, the PAs were 89.95% and 86.32%, respectively.

**Table 5.** Accuracy metrics of the *S. salsa* maps from random forest model (OA: overall accuracy; PA: producer’s accuracy; UA: user’s accuracy; Kappa: Kappa coefficient).

| Region/Year | Scheme | PA (%) |  | UA (%) |  | Kappa | OA (%) |
| --- | --- | --- | --- | --- | --- | --- | --- |
|  |  | SS | NSS | SS | NSS |  |  |
| YRD 2020 | With SSHI | 89.95 | 96.24 | 92.75 | 94.71 | 0.87 | **94.05** |
|  | W/O SSHI | 84.92 | 93.28 | 87.11 | 92.04 | 0.79 | 90.37 |
| YRD 2021 | With SSHI | 86.32 | 95.77 | 90.61 | 93.67 | 0.83 | **92.73** |
|  | W/O SSHI | 79.47 | 93.03 | 84.36 | 90.56 | 0.74 | 88.68 |
| YRD 2023 | With SSHI | 91.28 | 93.27 | 86.83 | 95.65 | 0.83 | **92.62** |
|  | W/O SSHI | 90.26 | 90.52 | 82.24 | 95.03 | 0.79 | 90.44 |
| LRD 2022 | With SSHI | 93.41 | 95.47 | 90.91 | 96.76 | 0.88 | **94.79** |
|  | W/O SSHI | 90.11 | 94.40 | 88.65 | 95.16 | 0.84 | 93.00 |
| LRD 2023 | With SSHI | 91.67 | 96.61 | 94.29 | 95.00 | 0.89 | **94.74** |
|  | W/O SSHI | 90.56 | 94.24 | 90.56 | 94.24 | 0.85 | 92.84 |

Figure 8 displays the top 10 features ranked by their SHAP values. Across all images in both study areas, SSHI consistently proves to be the most important feature with the highest SHAP value, highlighting its strong contribution to *S. salsa* mapping. Notably, SSHI’s SHAP value significantly exceeds those of other features, except in the 2023 YRD image (Figure 8c). This aligns with findings in Table 5, where excluding SSHI leads to reduced classification accuracy. Other key features include surface reflectance in wavebands between 530 nm and 570 nm, as well as NIR and SWIR bands.

**Figure 8.**
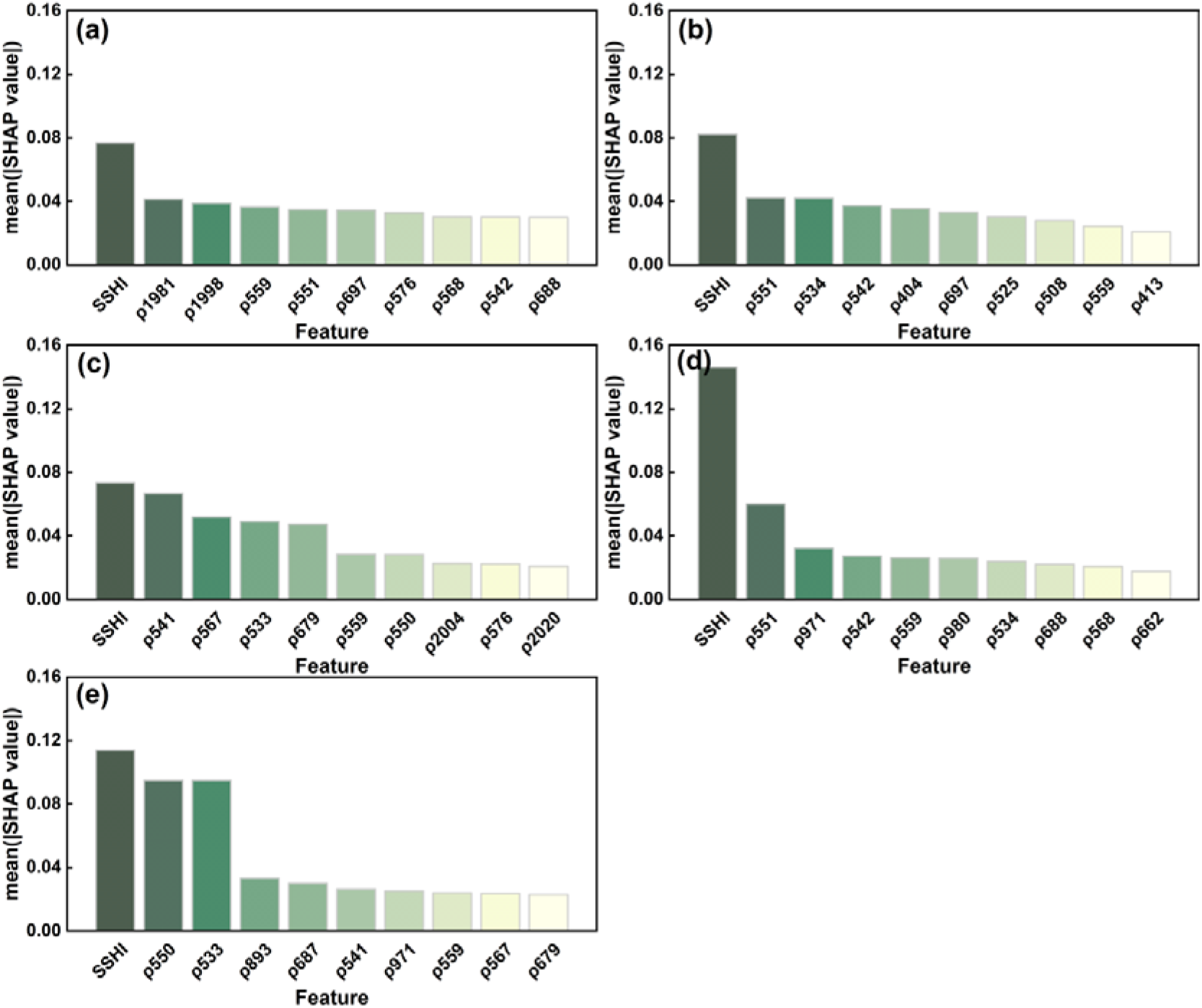
Mean SHAP values of the features with the highest 10 SHAP values for (a) YRD in 2020, (b) YRD in 2021, (c) YRD in 2023, (d) LRD in 2022, and (e) LRD in 2023.

Machine learning models generally produce higher accuracies than simple thresholding methods, as they can capture more complex relationships. Comparing Table 4 and Table 5 reveals that the RF models incorporating SSHI and surface reflectance achieved higher accuracies than the optimal thresholds. However, RF based solely on spectral reflectance provided only a slight accuracy improvement over thresholding methods. Both the SSHI thresholding and RF methods demonstrated the effectiveness and reliability of SSHI, showing that using only an SSHI threshold yielded good accuracy and significantly enhanced the RF classification results. Given that the thresholding method requires only SSHI as a classification feature while delivering reliable performance, the SSHI thresholding approach is efficient and has potential to be applied to large-scale mapping of *S. salsa*.

### 4.4 Analysis of *S. salsa* maps in different years

Figure 9 and Figure 10 show the *S. salsa* maps derived from the RF models with SSHI and surface reflectance at the 35 wavebands in YRD and LRD. In YRD (Figure 9), *S. salsa* was primarily distributed in the mid-tidal flats on both sides of the river banks, between *S. alterniflora* in the lower intertidal zones and *P. australis* near the river channels. The estimated area of *S. salsa* in 2020, 2021 and 2023 in YRD were 2185hm^2^, 1776hm^2^, and 2106hm^2^, respectively. Due to cloud cover in the 2020 images, the actual extent of *S. salsa* may exceed 2185 hm^2^.

**Figure 9.**
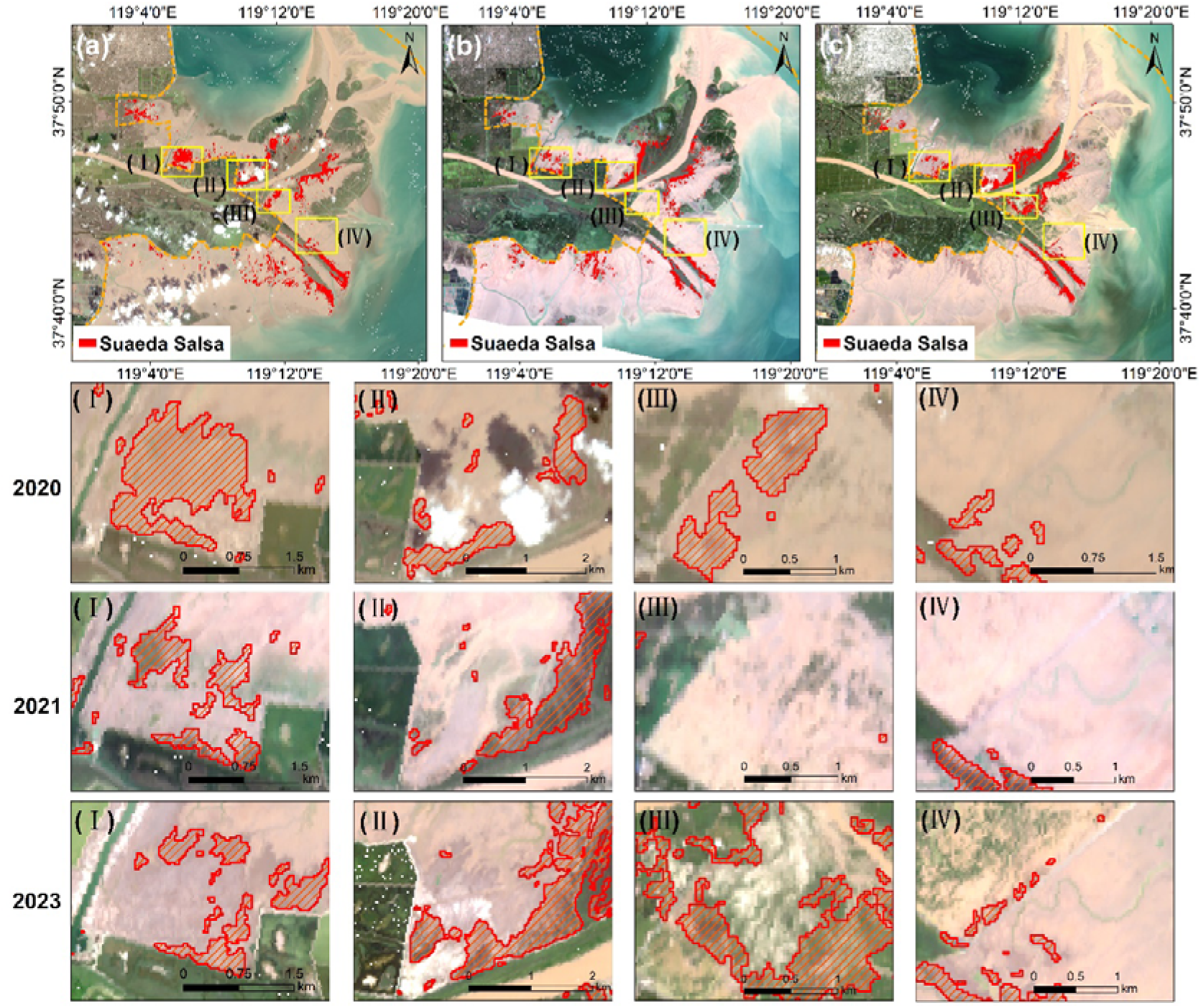
*S. salsa* maps in YRD for (a) 2020, (b) 2021, and (c) 2023; the second to fourth rows illustrate the zoom-in maps at subregions I, II, III and IV.

**Figure 10.**
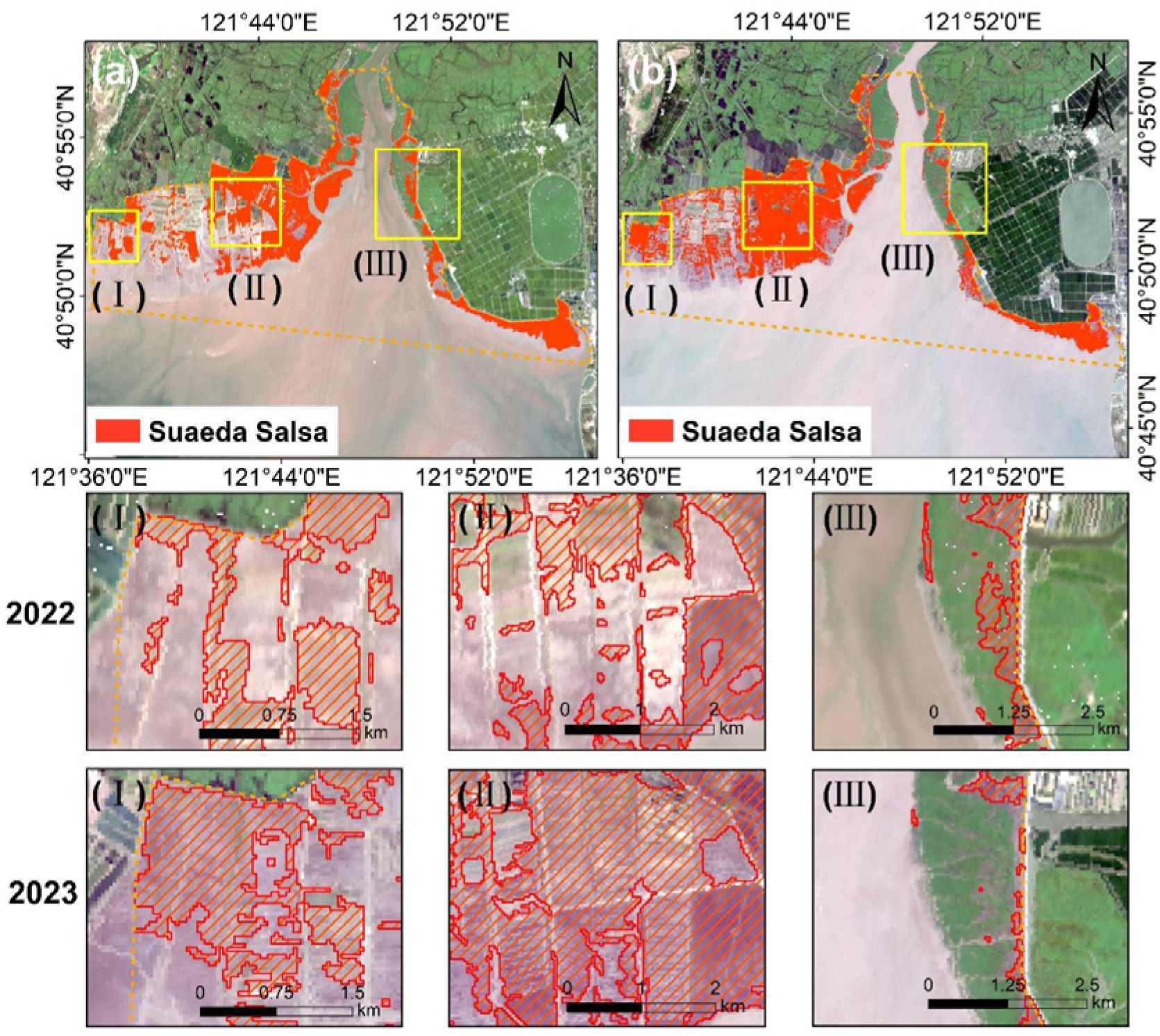
Mapping results of *S. salsa* in LRD for (a) 2022, and (b) 2023; (I), (II), and (III) highlight regions with interannual variation characteristics.

Subregions I-IV in Figure 9 reveal a decline in *S. salsa* from 2020 to 2021, followed by slight recovery in 2023, aligning with restoration efforts initiated in 2019 [49]. However, some restored patches were not well-preserved in 2021. Notably, Subregion III, which featured intact patches established by YRDNNR in 2020, witnessed a complete loss of these patches in 2021. This loss likely resulted from the severe flooding in the Yellow River Delta due to heavy rainfall in the lower Yellow River from August to October. In September, discharge rates around YRD exceeded 5,240 m³/s, submerging significant areas of tidal flats with heavy sediment, which damaged the restored patches. From late 2021 to 2022, large-scale *S. alterniflora* removal projects were conducted in YRD [23]. By 2023, no large *S. alterniflora* patches remained (Figure 9c). The removal of this invasive species may have improved hydrological connectivity on tidal flats, facilitating the recovery of *S. salsa* in 2023 [50].

In LRD, *S. salsa* dominated the tidal flats, with extensive and intact patches along the river banks. The area of *S. salsa* was 4310 hm^2^ in 2022 and 4348 hm^2^ in 2023, showing a slight increase [51]. This is mainly due to the continuous implementation of “returning aquaculture ponds to wetlands” and “returning farmland to wetlands” projects in LRD, which led to the conversion of large areas of agricultural land and aquaculture ponds into *S. salsa* from 2022 to 2023, proving the effectiveness of these projects (Figure10 (I)∼(II)). However, there are still some areas where the area of *S. salsa* has decreased. This may be due to *S. salsa*, as a pioneer species, accelerated sedimentation and reduced soil salinity, promoting *P. australis* colonization [52]. Rainfall reduces the salinity of the surface soil of tidal flat, making it more suitable for the growth of *P. australis*. Therefore, *P. australis* can outcompete*S. salsa* in later stages of succession, leading to its dominance in some areas (Figure 10(III)) [53]. At the same time, climate conditions may also be one of the factors affecting the change in the area of *S. salsa*. The climate in 2023 is relatively dry, with a precipitation of 437.6 mm, compared to 888.8 mm in 2022. The reduced soil moisture and increased salinity are unfavourable for *S. salsa* growth. Under the influence of hydrological conditions, some areas may have been submerged due to proximity to water, contributing to the identified decrease [5, 54].

## 5. Discussion

### 5.1 Characteristics of SSHI

The development of SSHI took into account the unique characteristics of both coastal salt marsh wetlands and hyperspectral imagery. Coastal wetlands are characterized by patchy vegetation distribution with significant variations in coverages, as well as substantial spatial and temporal fluctuations in soil moisture due to tidal influences, resulting in substantial within-class spectral variations. To address this, we employed a stratified sample selection approach and analyzed the spectral curves of the samples with varying vegetation coverages and soil moisture. This allowed us to identify feature bands capable of distinguishing *S. salsa*, irrespective of coverage variations. Second, we utilized multiple measurements, including correlation coefficient, OIF, and J-M distance, to determine optimal band combinations to avoid spectral redundancy inherent in hyperspectral data and to represent the optimal separation between different cover types.

The novelty of SSHI also resides in its adaptability to each pixel. Unlike previous hyperspectral indices based on several fixed bands, SSHI uses flexible band selection for each pixel, i.e., the central band where the peak or valley within a specified window was located. max(p_i_) ( i E )499nm,611nm] ) captures the peak reflectance in the green band, and min(p_j_) ( jE )628nm,851nm]) enhances the difference between *S. salsa* and bare flat. To further illustrate the advantages of the flexible wavebands, we tested a fixed green peak at 568nm and a fixed NIR reflectance at 671nm. The fixed-band index formula was: 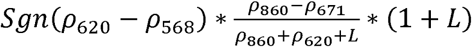.

Figure 11 shows considerable overlaps between the green vegetation and *S. salsa* for fixed-band SSHI (Figure 11b), in comparison with our SSHI, which can be attributed to the variability in the spectral peaks in the visible wavelengths for low-density green vegetation. In such cases, the difference in reflectance between 620nm and 568nm could produce positive values, leading to inaccuracies (Figure 12). By dynamically identifying local maximum and minimum reflectance values for each pixel (Figure 12), more accurate pixel-by-pixel analysis and classification can be achieved, effectively distinguishing *S. salsa* from other land cover types.

**Figure 11.**
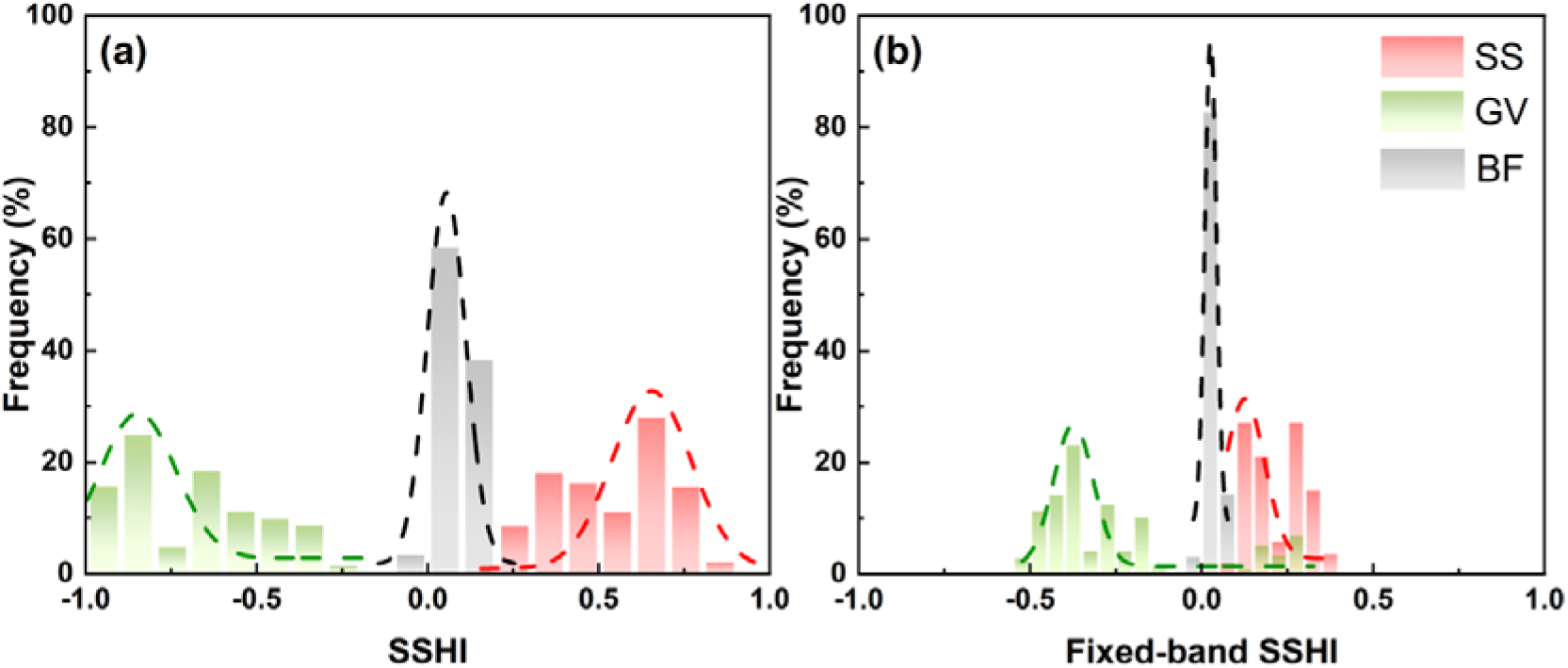
(a) SSHI histogram and (b) fixed-band SSHI histogram of *S. salsa*, green vegetation and bare flats. The samples in Section 3.1.1 were used for analysis.

**Figure 12.**
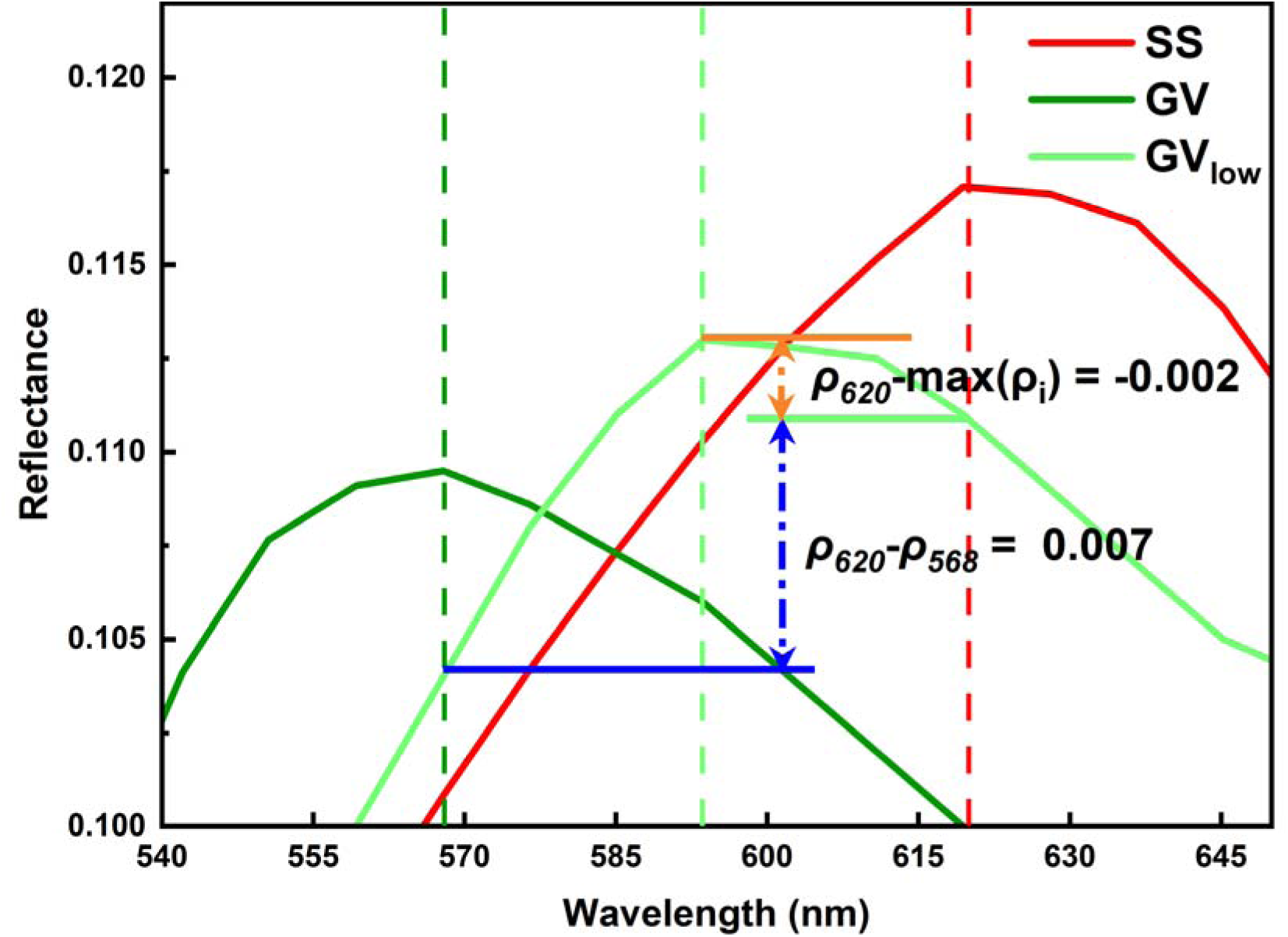
Schematic diagram of (p_620_ - p_S68_) which was used in fixed-band SSHI and p_620_ - max(p_l_), (i E [499nm, 611nm]) used in our SSHI for different land cover types. SS represents *S. salsa*, GV represents green vegetation, and GV_low_ represents a low-coverage green vegetation (Level 1). Low-coverage green vegetation has negative p_620_ - max(p_l_), while positive (p_620_ - p_S68_).

### 5.2 Applicability of SSHI to other hyperspectral images

To evaluate the applicability of SSHI on different hyperspectral images, we tested SSHI on GF-5B AHSI imagery and Hyperspectral Precursor of the Application Mission (PRISMA) hyperspectral imagery over YRD and LRD during the growing season. GF-5B satellite, launched by China on September 7, 2021, carries AHSI providing 30m-resolution imagery with 60 km swath width across 330 wavebands ranging from 400 to 2500nm, including 150 VNIR bands with 5nm spectral resolution and 180 SWIR bands with 10nm resolution. PRISMA hyperspectral system collects images with 30 km swath, 30 m spatial resolution and 239 wavebands (66 bands at VNIR and 173 bands at SWIR from 400∼2500nm) with spectral sampling interval < 12 nm.

Due to limited availability, only three scenes in 2023, including one GF-5B ASHI image on August 31 over YRD, one GF-5B ASHI image on August 10 over LRD and one PRISMA image on September 19 over LRD, were collected (Table 6). All images were preprocessed using the same procedures as ZY1-02D/E imagery (Section 2.2.1). We calculated GF-5B-based SSHI and PRISMA-based SSHI for the three images using the central wavebands closest to the ZY-02D AHSI bands, i.e.,

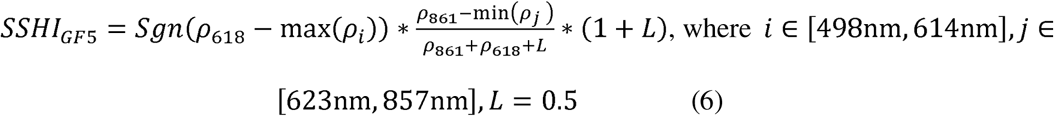

**Table 6.** Accuracy metrics of the *S. salsa* maps based on GF-5B AHSI and PRISMA imagery. OA: overall accuracy; PA: producer’s accuracy; UA: user’s accuracy; Kappa: Kappa coefficient; Thres: threshold.

| Image/Date | Region | Method | PA (%) |  | UA (%) |  | Kappa | OA (%) |
| --- | --- | --- | --- | --- | --- | --- | --- | --- |
|  |  |  | SS | NSS | SS | NSS |  |  |
| GF-5B AHSI<br>2023-08-31 | YRD | Thres=0.24 | 80.00 | 90.82 | 80.83 | 90.37 | 0.71 | **87.29** |
|  |  | RF | 88.21 | 80.37 | 89.58 | 94.01 | 0.76 | **89.13** |
| GF-5B AHSI<br>2023-08-10 | LRD | Thres=0.19 | 81.82 | 95.32 | 91.72 | 89.23 | 0.79 | **90.09** |
|  |  | RF | 87.50 | 95.68 | 92.77 | 92.36 | 0.84 | **92.51** |
| PRISMA<br>2023-09-19 | LRD | Thres=0.20 | 84.44 | 88.81 | 82.16 | 90.34 | 0.73 | **87.16** |
|  |  | RF | 83.89 | 96.62 | 93.79 | 90.79 | 0.82 | **91.81** |

and,

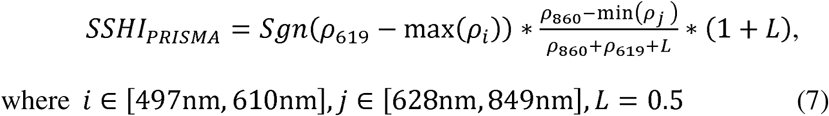

where SSHI_GF5_ and SSHI_PRIsMA_ denote SSHI for GF-5B and PRISMA imagery, respectively.

We utilized both thresholding and RF (with SSHI) methods using the same sample sets for ZY1-02D/E imagery in the same year. Table 6 shows that the OAs of GF-5B thresholding-derived *S. salsa* maps are 87.29% in YRD and 90.09% in LRD; the OAs of RF-derived maps are higher, i.e., 89.13% in YRD and 92.51% in LRD. The OAs from PRISMA imagery are 87.16% for thresholding method and 91.81% for RF method. For both satellite imagery, the resultant *S. salsa* maps have reasonable accuracies, although they are slightly lower than those from ZY1-02D/E imagery. In addition, the thresholds for both imageries are similar as those for ZY1-02D/E imagery, suggesting robustness of SSHI for different hyperspectral imagery.

### 5.3 Potential and implications

Our results indicate that SSHI demonstrated good capability to identify *S. salsa* and significantly improves the accuracy of *S. salsa* extraction, particularly in YRD where coverage is low. Compared to existing red pigment indices, SSHI provides better separation of *S. salsa* from other land cover types. Moreover, SSHI demonstrates robustness, achieving high accuracy at both study areas using different hyperspectral imagery. In recent years, China has been actively conducting ecological restoration projects for coastal wetlands. As a pioneer species, *S. salsa* is the major restoration target species. Our results suggest that SSHI offers a reliable method for monitoring *S. salsa* distribution. Additionally, SSHI, adopting the form of SAVI, offers potential for estimating *S. salsa* coverages and biomass, which are important indicators for evaluation of restoration effectiveness. Within the *S. salsa* extents at both YRD and LRD, SSHI exhibits good linear relationship with NDVI (Figure S2), which has been widely recognized as indicator for coverage and biomass estimation. Our future research would further explore SSHI’s capability to estimate coverage and biomass. We believe that SSHI has great potential to provide a scientific method for evaluation of *S. salsa* restoration effectiveness.

The *S. salsa* maps reveal that the species is vulnerable to hydrological and climate change. The removal of invasive *S. alterniflora* in YRD may alter the hydrological connectivity at the tidal flats, which may further alter the distribution and growth of *S. salsa* [49]. Severe river floods may damage *S. salsa* patches. Decreased soil water content and excessive soil salinity due to drought also led to *S. salsa* loss in LRD [4, 10]. These issues offer opportunities for the application of SSHI in in-depth studies on the mechanism of *S. salsa* recovery and degradation at coastal wetlands.

### 5.4 Uncertainties and limitations

At tidal wetlands in North China, plant species such as *S. salsa*, *S. alterniflora* and *P. australis* generally exist as mono-dominated or mosaic communities, forming the foundation of successful *S. salsa* identification at both study areas. However, at the ecotones of *S. salsa* and other species, mixed pixel issues can be inevitable, which may lead to misclassifications at these areas. These issues may be addressed by improving spatial resolution of the hyperspectral imagery via spectral-spatial fusion methods [55, 56] or utilization of high-spatial-resolution hyperspectral images.

Second, *S. salsa* generally exhibits red tones at coastal North China during the entire growing season because of the low temperature and high salinity [4, 10]. Therefore, SSHI can successfully distinguish *S. salsa* and green vegetations. However, in eastern coastal China such as Yellow Sea wetlands, *S. salsa* on tidal flats with lower soil salinity show green tones during spring and summer, and turn red in autumn. In these areas, SSHI-based *S. salsa* extraction during spring and summer could cause confusions. Therefore, appropriate selection of image acquisition date is critical for accurate extraction of *S. salsa* for these areas. Tidal level is another consideration factor for image selection. Extremely high tidal level may cause submergence of *S. salsa*, resulting in large omission errors. Therefore, it is highly suggested that tidal levels at the time of image acquisition should be examined.

## 6. Conclusion

In this study, we introduced a hyperspectral index named SSHI for *S. salsa* identification. The SSHI leverages the spectral variations in bare flats and vegetation at coastal wetlands, employing dynamic band selection on a per-pixel basis to optimize the separation of *S. salsa* from other land cover types. At both YRD and LRD, the SSHI, derived from ZY1-02D/E imagery, effectively identified *S. salsa* using either thresholding or RF methods, achieving high overall accuracies (OA=87.96%∼94.79%). While RF with SSHI and spectral reflectance marginally outperformed SSHI-based thresholding, the SSHI significantly enhanced classification accuracies, with PA and UA increasing by 1.02%∼6.85% compared to methods without SSHI. Compared to existing red-pigment-related hyperspectral indices, SSHI demonstrated superior separation of *S. salsa* from other land covers. Its performance was robust across various hyperspectral satellite imagery, including GF-5B and PRISMA, with thresholds consistent with those derived from ZY1-02D/E imagery. These findings highlight SSHI’s potential as a reliable and efficient tool for evaluating *S. salsa* restoration. Future studies could focus on quantitatively estimating *S. salsa* biophysical parameters using SSHI.

## CRediT authorship contribution statement

**Mengyao Zhang:** Methodology, Writing – original draft. **Yinghai Ke:** Methodology, Writing – original draft, Writing – review & editing. **Han Liu:** Writing – review & editing. **Zhaojun Zhuo:** Investigation, Methodology. **Kun Shang:** Writing – review & editing. **Peng Li:** Writing – review & editing. **Nana Zhao:** Writing – review & editing. **Jinghan Sha:** Investigation, Methodology. **Jinyuan Li:** Investigation, Methodology. **Yue Han:** Investigation, Methodology.

## Acknowledgments

The work is supported by National Natural Science Foundation of China (42071396, 42301461).

## Data availability

Data will be made available on request.

## References

[1] Song, Z., Sun, Y., Chen, P., and Jia, M. Assessing the ecosystem health of coastal wetland vegetation (Suaeda salsa) using the pressure state response model, a case of the Liao River estuary in China. International Journal of Environmental Research and Public Health. 2022;19(1):546 10.3390/ijerph19010546

[2] Gu, J., Jin, R., Chen, G., Ye, Z., Li, Q., Wang, H., Li, D., Christakos, G., Agusti, S., and Duarte, C.M. Areal extent, species composition, and spatial distribution of coastal saltmarshes in China. IEEE Journal of Selected Topics in Applied Earth Observations and Remote Sensing. 2021;147085–7094 10.1109/JSTARS.2021.3093673

[3] Li, M., Zhao, Z., Liu, J., Ma, S., and He, P. Research advance on the synthesis and regulation mechanism of betacyanins in Suaeda salsa. Plant Physiology Journal. 2023;60(01):63–74 10.13592/j.cnki.ppj.300121.

[4] Zhang, S., Tian, J., Lu, X., Tian, Q., He, S., Lin, Y., Li, S., Zheng, W., Wen, T., and Mu, X. Monitoring of chlorophyll content in local saltwort species Suaeda salsa under water and salt stress based on the PROSAIL-D model in coastal wetland. Remote Sensing of Environment. 2024;306114117 10.1016/j.rse.2024.114117

[5] Cao, C., Su, F., Song, F., Yan, H., and Pang, Q. Distribution and disturbance dynamics of habitats suitable for Suaeda salsa. Ecological Indicators. 2022;140108984 10.1016/j.ecolind.2022.108984

[6] Debanshi, S., and Pal, S. Modelling water richness and habitat suitability of the wetlands and measuring their spatial linkages in mature Ganges delta of India. Journal of Environmental Management. 2020;271110956 10.1016/j.jenvman.2020.110956

[7] Zhang, C., Gong, Z., Qiu, H., Zhang, Y., and Zhou, D. Mapping typical salt-marsh species in the Yellow River Delta wetland supported by temporal-spatial-spectral multidimensional features. Science of The Total Environment. 2021;783147061 10.1016/j.scitotenv.2021.147061

[8] Ahmed, K.R., Akter, S., Marandi, A., and Schüth, C. A simple and robust wetland classification approach by using optical indices, unsupervised and supervised machine learning algorithms. Remote Sensing Applications: Society and Environment. 2021;23100569 10.1016/j.rsase.2021.100569

[9] Fan, P., Cai, Y., Liu, J., Wu, N., and Guo, X. Monitoring Suaeda Salsa Changes in the Yellow River Delta Based on Landsat Satellite Images. Geomatics & Spatial Information Technology. 2023;46(03):38–41 10.3969/j.issn.1672-5867.2023.03.012

[10] Ke, Y., Han, Y., Cui, L., Sun, P., Min, Y., Wang, Z., Zhuo, Z., Zhou, Q., Yin, X., and Zhou, D. Suaeda salsa spectral index for Suaeda salsa mapping and fractional cover estimation in intertidal wetlands. ISPRS Journal of Photogrammetry and Remote Sensing. 2024;207104–121 10.1016/j.isprsjprs.2023.11.018

[11] Zhang, H., Hu, M., Ma, H., Jiang, L., Zhao, Z., Ma, J., and Wang, L. Differential responses of dimorphic seeds and seedlings to abiotic stresses in the halophyte Suaeda salsa. Frontiers in Plant Science. 2021;12630338 10.3389/fpls.2021.630338

[12] Wang, Z., Ke, Y., Lu, D., Zhuo, Z., Zhou, Q., Han, Y., Sun, P., Gong, Z., and Zhou, D. Estimating fractional cover of saltmarsh vegetation species in coastal wetlands in the Yellow River Delta, China using ensemble learning model. Frontiers in Marine Science. 2022;91077907 10.3389/fmars.2022.1077907

[13] Wang, Z., Ke, Y., Chen, M., Zhou, D., Zhu, L., and Bai, J. Mapping coastal wetlands in the Yellow River Delta, China during 2008–2019: impacts of valid observations, harmonic regression, and critical months. International Journal of Remote Sensing. 2021;42(20):7880–7906 10.1080/01431161.2021.1966852

[14] Han, Y., Ke, Y., Wang, Z., Liang, D., and Zhou, D. Classification of the Yellow River Delta wetland landscape based on ZY1-02D hyperspectral imagery. National Remote Sensing Bulletin. 2023;27(06):1387–1399 10.11834/jrs.20211071

[15] Cheng, S., Yang, X., Yang, G., Chen, B., Chen, D., Wang, J., Ren, K., and Sun, W. Using ZY1-02D satellite hyperspectral remote sensing to monitor landscape diversity and its spatial scaling change in the Yellow River Estuary. International Journal of Applied Earth Observation and Geoinformation. 2024;128103716 10.1016/j.jag.2024.103716

[16] Guo, S., Feng, Z., Wang, P., Chang, J., Han, H., Li, H., Chen, C., and Du, W. Mapping and Classification of the Liaohe Estuary Wetland Based on the Combination of Object-Oriented and Temporal Features. IEEE Access. 2024;10.1109/ACCESS.2024.3389935

[17] Sun, W., Liu, K., Ren, G., Liu, W., Yang, G., Meng, X., and Peng, J. A simple and effective spectral-spatial method for mapping large-scale coastal wetlands using China ZY1-02D satellite hyperspectral images. International Journal of Applied Earth Observation and Geoinformation. 2021;104102572 10.1016/j.jag.2021.102572

[18] Yang, G., Huang, K., Sun, W., Meng, X., Mao, D., and Ge, Y. Enhanced mangrove vegetation index based on hyperspectral images for mapping mangrove. ISPRS Journal of Photogrammetry and Remote Sensing. 2022;189236–254 10.1016/j.isprsjprs.2022.05.003

[19] Yang, D., Chen, J., Zhou, Y., Chen, X., Chen, X., and Cao, X. Mapping plastic greenhouse with medium spatial resolution satellite data: Development of a new spectral index. ISPRS Journal of Photogrammetry and Remote Sensing. 2017;12847–60 10.1016/j.isprsjprs.2017.03.002

[20] Yue, X., Sun, Q., Lv, M., Li, J., Wang, A., Zhang, S., Xie, T., and Wang, Q. Evaluation of Effects of Different Restoration Measures on Suaeda Salsa Wetland in Yellow River Delta. EnEng. 2024;1–13

[21] Wang, X., Xiao, X., Zhang, X., Wu, J., and Li, B. Rapid and large changes in coastal wetland structure in China’s four major river deltas. Global Change Biology. 2023;29(8):2286–2300 10.1111/gcb.16583

[22] Tan, K., Sun, D., Dou, W., Wang, B., Sun, Q., Liu, X., Zhang, H., Lan, Y., and Lun, F. Mapping Coastal Wetlands and Their Dynamics in the Yellow River Delta over Last Three Decades: Based on a Spectral Endmember Space. Remote Sensing. 2023;15(20):5003 10.3390/rs15205003

[23] Min, Y., Cui, L., Li, J., Han, Y., Zhuo, Z., Yin, X., Zhou, D., and Ke, Y.J.I.J.o.A.E.O. Detection of large-scale Spartina alterniflora removal in coastal wetlands based on Sentinel-2 and Landsat 8 imagery on Google Earth Engine. International Journal of Applied Earth Observation and Geoinformation. 2023;125103567 10.1016/j.jag.2023.103567

[24] Wang, W. Remote sensing monitoring research on the succession and degradation of typical Saline vegetation community. Dalian Ocean University, 2022. 10.27821/d.cnki.gdlhy.2022.000043

[25] Liu, S., Chen, H., Xing, Q., Cheng, H., Han, J., and XU, X. Ecosystem CO_2_ Exchange and Its Environmental Regulation of a Restored Wetland in the Liaohe River Estuary. Environmental Science. 2024;45(02):920–928 10.13227/j.hjkx.202303247.

[26] Hu, J. Repairing saline alkali land and protecting the Red Beach. People’s Daily. 014. 2024, 01–09. 10.28655/n.cnki.nrmrb.2024.000294.

[27] Shao, C., Yang, G., Sun, W., Zuo, Y., Ge, W., and Yang, S. Construction method of a Spartina alterniflora index based on hyperspectral satellite images. National Remote Sensing Bulletin. 2024;28(03):635–648 10.11834/jrs.20242621

[28] Zheng, S., Hai, Y., He, M., and Wang, J. Comparative study on vegetation extraction effect based on ZY1-02D data. Spacecraft Recovery & Remote Sensing. 2022;43(02):92–103 10.3969/j.issn.1009-8518.2022.02.010

[29] Awasthi, M.P. Mapping and analyzing temporal variability of spectral indices in the lowland region of Far Western Nepal. Water Practice and Technology. 2023;18(11):2971–2988 10.2166/wpt.2023.180

[30] Tahir, H., and Din, A.H.M. The Potential of Landsat 8 OLI Images in Coastline Identification: The Case Study of Basra, Iraq. *Engineering*, Technology and Applied Science Research. 2024;14(1):13041–13046 10.48084/etasr.6580

[31] Fu, Y. Changes of the vegetation and water and their responses to climatic variables in the intersection area of multi water resources in western Jinan (master), University of Jinan, 2023. 10.27166/d.cnki.gsdcc.2023.001319.

[32] Sun, X., Hou, D., and Bai, y. Research on Soil Moisture Inversion Based on Sentinel-1 and Landsat8-OLI Data. Gansu Water Resources and Hydropower. 2024;60(01):29–32 10.19645/j.issn2095-0144.2024.01.007.

[33] Sun, W., and Du, Q. Hyperspectral band selection: A review. IEEE Geoscience and Remote Sensing Magazine. 2019;7(2):118–139 10.1109/MGRS.2019.2911100

[34] Figliomeni, F.G., Guastaferro, F., Parente, C., and Vallario, A. A Proposal for Automatic Coastline Extraction from Landsat 8 OLI Images Combining Modified Optimum Index Factor (MOIF) and K-Means. Remote Sensing. 2023;15(12):3181 10.3390/rs15123181

[35] Kong, Y., Wang, L., Feng, H., Xu, Y., Liang, L., Xu, L., Yang, X., and Zhang, Q. Leaf area index estimation based on UAV hyperspectral band selection. Spectroscopy and Spectral Analysis. 2022;42(03):933–939 10.3964/j.issn.1000-0593(2022)03-0933-07

[36] Van, N.T.G., McVicar, T.R., and Datt, B. On the relationship between training sample size and data dimensionality: Monte Carlo analysis of broadband multi-temporal classification. Remote sensing of environment. 2005;98(4):468–480 10.1016/j.rse.2005.08.011

[37] Zhang, X., Lin, X., Fu, D., Wang, Y., Sun, S., Wang, F., Wang, C., Xiao, Z., and Shi, Y. Comparison of the Applicability of JM Distance Feature Selection Methods for Coastal Wetland Classification. Water. 2023;15(12):2212 10.3390/w15122212

[38] Huete, A.R. A soil-adjusted vegetation index (SAVI). Remote sensing of environment. 1988;25(3):295–309 10.1016/0034-4257(88)90106-x

[39] Breiman, L. Random forests. Machine learning. 2001;455–32 10.1023/A:1010933404324

[40] Jiao, L., Sun, W., Yang, G., Ren, G., and Liu, Y. A hierarchical classification framework of satellite multispectral/hyperspectral images for mapping coastal wetlands. Remote Sensing. 2019;11(19):2238

41. [41] Lundberg, S. A unified approach to interpreting model predictions. *arXiv preprint arXiv:.07874.* 2017;

[42] Sakuta, M. Diversity in plant red pigments: anthocyanins and betacyanins. Plant biotechnology reports. 2014;837–48 10.1007/s11816-013-0294-z

[43] Meyer, G.E., and Neto, J.C. Verification of color vegetation indices for automated crop imaging applications. Computers and electronics in agriculture. 2008;63(2):282–293 10.1016/j.compag.2008.03.009

[44] Gamon, J., and Surfus, J. Assessing leaf pigment content and activity with a reflectometer. The New Phytologist. 1999;143(1):105–117 10.1046/j.1469-8137.1999.00424.x

[45] Gitelson, A.A., Merzlyak, M.N., and Chivkunova, O.B. Optical properties and nondestructive estimation of anthocyanin content in plant leaves. Photochemistry and photobiology. 2001;74(1):38–45 10.1562/0031-8655(2001)0740038OPANEO2.0.CO2

[46] Li, Y., and Huang, J. Leaf anthocyanin content retrieval with partial least squares and gaussian process regression from spectral reflectance data. Sensors. 2021;21(9):3078 10.3390/s21093078

[47] Somers, B., and Asner, G.P. Multi-temporal hyperspectral mixture analysis and feature selection for invasive species mapping in rainforests. Remote Sensing of Environment. 2013;13614–27 10.1016/j.rse.2013.04.006

[48] Congalton, R.G. A comparison of sampling schemes used in generating error matrices for assessing the accuracy of maps generated from remotely sensed data. Photogrammetric Engineering and Remote Sensing. 1988;54593–600

[49] Ning, Z., Chen, C., Xie, T., Li, S., Zhu, Z., Wang, Q., Cai, Y., Bai, J., and Cui, B. Invasive plant indirectly affects its self-expansion and native species via bio-geomorphic feedbacks: Implications for salt marsh restoration. Catena. 2023;226107056 10.1016/j.catena.2023.107056

[50] Ren, G., Zhao, Y., Wang, J., Wu, P., and Ma, Y. Ecological effects analysis of Spartina alterniflora invasion within Yellow River delta using long time series remote sensing imagery. Estuarine, Coastal and Shelf Science. 2021;249107111 10.1016/j.ecss.2020.107111

[51] Li, Y., Wang, Z., Zhao, C., Jia, M., Ren, C., Mao, D., and Yu, H. Research on spatial-temporal dynamics of Suaeda salsa in Liaohe estuary and its identification mechanism using remote sensing. Remote Sensing for Natural Resources. 2024;1–9 10.1759.P.20240731.1443.006.html.

[52] Zhang, M. Research advancement on degradation mechanism and ecological restoration technology of coastal salt-marsh: a review. Journal of Dalian Ocean University. 2022;37(04):539–549 10.16535/j.cnki.dlhyxb.2022-181.

[53] Chen, X., Zhang, M., and Zhang, W. Landscape pattern changes and its drivers inferred from salt marsh plant variations in the coastal wetlands of the Liao River Estuary, China. Ecological Indicators. 2022;145109719 10.1016/j.ecolind.2022.109719

[54] Yu, Z., Yin, S., Bai, J., Wang, C., Chen, G., Wang, W., Wang, Y., Cui, B., Liu, X., and Li, X. Suaeda salsa in Relation to Hydrological Connectivity: From the View of Plant Trait Networks. Journal of Environmental Informatics. 2024;43(1):10.3808/jei.202400507

[55] Chen, N., Sui, L., Zhang, B., He, H., Gao, K., Li, Y., Junior, J.M., and Li, J. Fusion of Hyperspectral-Multispectral images joining Spatial-Spectral Dual-Dictionary and structured sparse Low-rank representation. International Journal of Applied Earth Observation and Geoinformation. 2021;104102570 10.1016/j.jag.2021.102570

[56] Zhang, M., Zheng, G., Jiang, Z., Zhu, Q., Wang, L., and Guan, Q. Local-aware coupled network for hyperspectral image super-resolution. GIScience and Remote Sensing. 2023;60(1):2233725 10.1080/15481603.2023.2233725

