## Supplemental Table S1, Figure S1 and S2 for "Suaeda Salsa Hyperspectral Index (SSHI) for mapping *S. salsa* in coastal wetlands using hyperspectral satellite imagery"

**Contents of this file**

Tables S1

Figures S1 to S2

**Table S1.** Examples of UAV image and Jilin-1 image within ZY1-02D AHSI pixel size (30m$\times$30m), the spectral reflectance curve of the corresponding ZY1-02D AHSI pixel and the calculated NDVI, and the corresponding class. SS-Level 1 to SS-Level 4 represent S. salsa with coverages from low to high; GV-Level 1 to GV -Level 4 represent green vegetation with coverages from low to high.

| UAV  (30m$\times$30m) | Jilin-1  (30m$\times$30m) | Spectral curve  (ZY1-02D AHSI) | NDVI (ZY1-02D AHSI) | Class |
| --- | --- | --- | --- | --- |
| 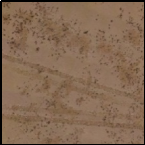 | 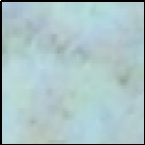 | 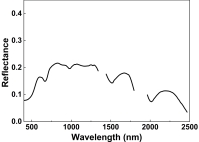 | 0.157 | SS-Level 1 |
| 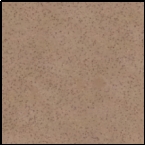 | 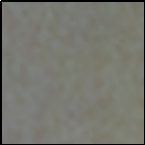 | 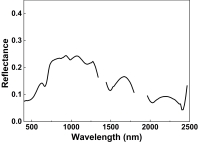 | 0.271 | SS-Level 2 |
| 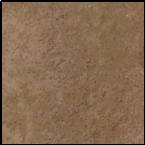 | 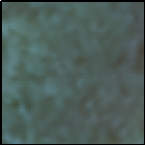 | 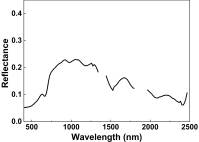 | 0.410 | SS-Level 3 |
| 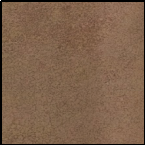 | 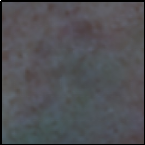 | 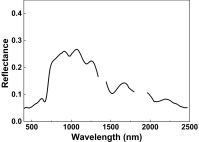 | 0.479 | SS-Level 4 |
| 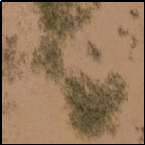 | 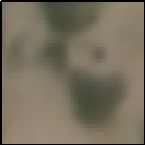 | 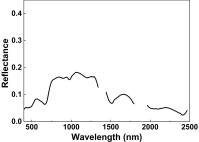 | 0.234 | GV-Level 1 |
| 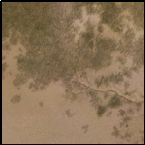 | 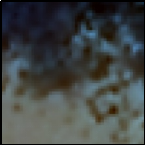 | 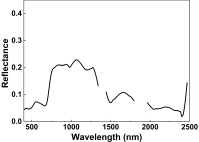 | 0.387 | GV-Level 2 |
| 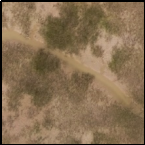 | 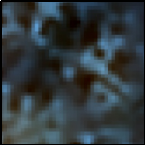 | 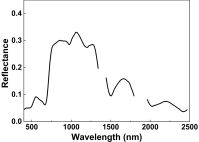 | 0.539 | GV-Level 3 |
| 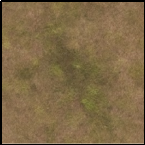 | 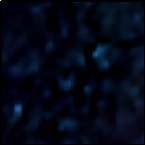 | 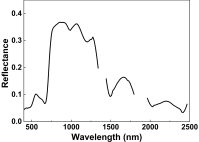 | 0.659 | GV-Level 4 |

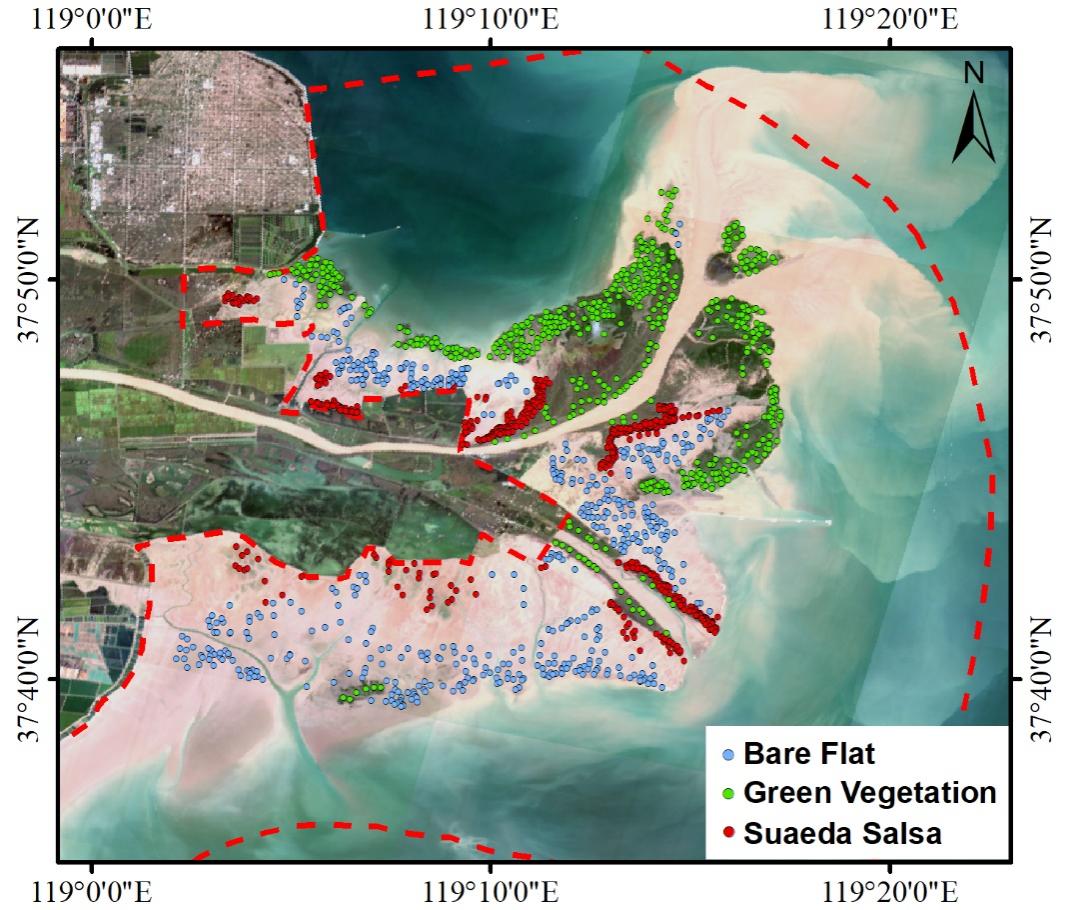

**Figure S2.** Location of the 2100 samples.

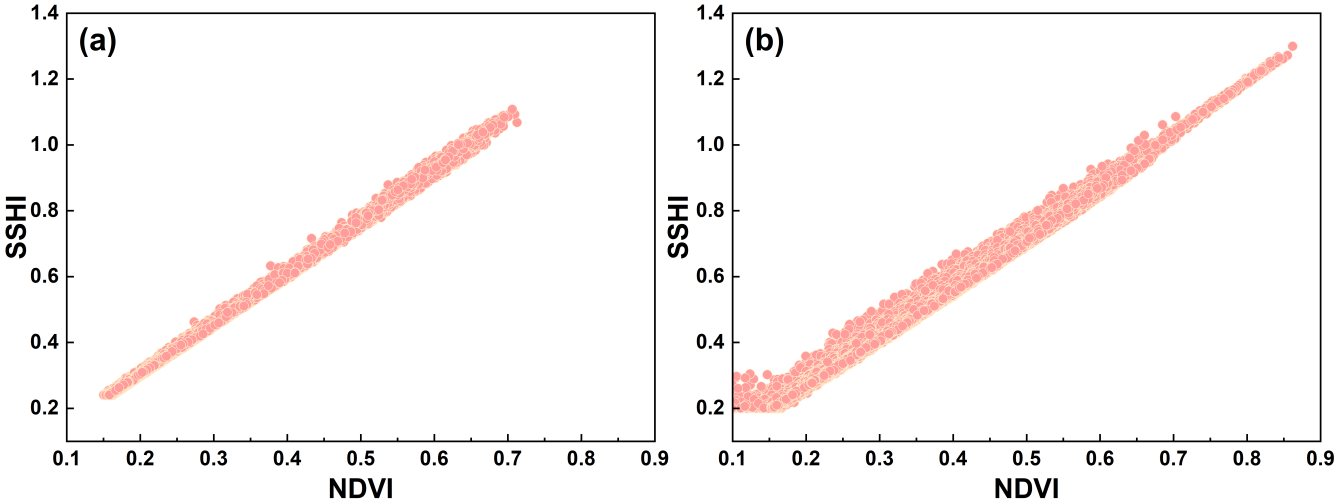

**Figure S3.** Scatter plot of SSHI and NDVI for *S. salsa* in (a) YRD and (b) LRD.
